# Uncovering smooth structures in single-cell data with PCS-guided neighbor embeddings

**DOI:** 10.1101/2025.06.27.661958

**Authors:** Rong Ma, Xi Li, Jingyuan Hu, Bin Yu

## Abstract

Single-cell sequencing is revolutionizing biology by enabling detailed investigations of cell-state transitions. Many biological processes unfold along continuous trajectories, yet it remains challenging to extract smooth, low-dimensional representations from inherently noisy, highdimensional single-cell data. Neighbor embedding (NE) algorithms, such as t-SNE and UMAP, are widely used to embed high-dimensional single-cell data into low dimensions. But they often introduce undesirable distortions, resulting in misleading interpretations. Existing evaluation methods for NE algorithms primarily focus on separating discrete cell types rather than capturing continuous cell-state transitions, while dynamic modeling approaches rely on strong assumptions about cellular processes and specialized data. To address these challenges, we build on the Predictability–Computability–Stability (PCS) framework for reliable and reproducible data-driven discoveries. First, we systematically evaluate popular NE algorithms through empirical analysis, simulation, and theory, and reveal their key shortcomings such as artifacts and instability. We then introduce NESS, a principled and interpretable machine learning approach to improve NE representations by leveraging algorithmic stability and to enable robust inference of smooth biological structures. NESS offers useful concepts, quantitative stability metrics, and efficient computational workflows to uncover developmental trajectories and cell-state transitions in single-cell data. Finally, we apply NESS to six single-cell datasets, including those about pluripotent stem cell differentiation, organoid development, and multiple tissue-specific lineage trajectories. Across these diverse contexts, NESS consistently yields useful and verifiable biological insights, such as identification of transitional and stable cell states and quantification of transcriptional dynamics during development. Notably, NESS resolves distinct neuronal subpopulations during embryoid formation and provides a deeper understanding of their cell-state dynamics.

## 1 Introduction

Recent advances in single-cell sequencing technologies have opened unprecedented opportunities to unravel cell-state transitions and their role in human health and disease. These technologies have facilitated the generation of high-resolution datasets that offer a detailed view of gene expression at the cellular level, shedding light on complex biological processes such as differentiation, reprogramming, and disease progression [83, 72, 66, 18, 63, 76, 24]. However, these datasets are inherently high-dimensional, sparse, and noisy due to technical limitations such as dropout events and measurement noise, making it challenging to extract biologically meaningful signals. Although single-cell data are often represented as sparse count matrices, the underlying cellular states typically reflect structured, continuous biological processes rather than discrete states. In many cases, these large-scale single-cell datasets exhibit lower-dimensional smooth manifold structures embedded within the original high-dimensional feature space, representing the dynamics of underlying biological processes. Examples of such low-dimensional smooth manifolds include linear trajectories [63, 28], cycles [45], and tree structures [34, 74]. Finding such structures and understanding their geometric properties can lead to important biological insights and knowledge [83, 72, 66, 18]. In particular, trajectory inference based on single-cell RNA sequencing (scRNA-seq) data aims to organize cells along a continuous trajectory in a lower-dimensional space, capturing the progression of biological processes [63, 76, 24]. And understanding characteristics of the manifold geometry, such as curvature, branching, and continuity, has offered valuable insights into the dynamics of cellular activity [66]. These works highlight the importance of reliable low-dimensional smooth representations for studying dynamic cellular processes.

A key challenge in analyzing single-cell omics data is to accurately represent and interpret biologically meaningful manifold structures in a low-dimensional space.^1^ To address this, neighbor embedding (NE) methods (such as t-SNE [71], UMAP [54], PHATE [55] and their extensions [69, 47, 56]) have been developed and become widely used tools for embedding high-dimensional single-cell data into more tractable low-dimensional representations. These methods typically begin by constructing an affinity graph among cells, followed by iterative updates to the low-dimensional embeddings that optimize a global objective function aimed at preserving local neighborhood relationships. Although originally developed for visualization, NE methods now play a central role in many state-of-the-art algorithms for single-cell analysis. For instance, many trajectory inference algorithms (e.g., PAGA [80], Monocle3 [16], VIA [67], Palantir [65], and CellRank [43, 79]) rely on the low-dimensional cell embeddings generated by NE methods as the foundation for reconstructing dynamic biological processes. RNA velocity analysis, which models transcriptional dynamics as smooth vector fields to predict future cell states, depends similarly on NE embeddings [41, 10, 26]. Tools such as Velocyto [41] and scVelo [9] project estimated future cell states into NE-derived low-dimensional spaces to visualize dynamic processes such as cellular differentiation, making NE methods a critical component of RNA velocity interpretation.

Despite their widespread use in many applications, the reliability of NE algorithms remains an open question. That is, whether these algorithms faithfully^2^ capture underlying smooth structures and related biological signals. While simplifications are sometimes necessary to extract meaningful biological insights from complex data[60], a key limitation of NE methods is their tendency to introduce substantial artifacts that can misrepresent underlying biological structures. This problem is exacerbated by the high-dimensional, noisy nature of single-cell data and the intrinsic complexity of biological processes, making faithful low-dimensional representation particularly challenging. Recent studies [38, 26, 17, 52, 50, 68] have highlighted that NE methods can generate misleading structures, such as artificial separation or clustering of cell populations, which do not reflect true biological structures. These artifacts can have significant downstream consequences, including misclassification of cell types [50, 17], spurious lineage relationships [68, 52], and incorrect inferences about differentiation trajectories [26]. Moreover, the results of NE algorithms can be highly sensitive to parameter tuning and distance metrics in the NE algorithms, with small changes in hyperparameters potentially leading to significant alterations in the inferred structures, affecting downstream biological conclusions [14, 32, 81, 38].

To address these limitations, several methods have been developed to assess the reliability of NE algorithms for a given dataset. For example, EMBEDR [37] and scDEED [81] generate random null data through permutation or re-sampling techniques to calculate a reliability score for every cell embedding based on the similarity between the cell’s 2D-embedding neighbors and pre-embedding neighbors. Opt-SNE [8] utilizes Kullback-Leibler divergence evaluation in real time to tailor the NE algorithm in a dataset-specific manner, enabling fast computation and hyperparameter selection to improve visualization of cluster structures. DynamicViz [68] creates bootstrap samples from the original data to generate a sequence of dynamic visualizations, and LOO [49] inspects the landscape of the objective function of specific NE optimization, each capturing the variability of the cell embedding against certain sample perturbations. While these methods are useful for detecting dubious cell embeddings^3^ and optimize hyperparameters of NE algorithms, they primarily focus on datasets with clear cluster structures, such as atlas-level single-cell datasets comprising distinct cell types, with the goal of better distinguishing these clusters. However, for applications involving smooth structures such as cell differentiation with continuously transitioning cell states, these methods typically do not account for the dynamic nature of the underlying biological processes. Consequently, their evaluation metrics often do not adequately assess the biological relevance of the reconstructed low-dimensional cell-state manifolds and provide only limited insight into the dynamics of cell-state transitions (see Results, Figures 4 and S10a). On the other hand, some recent works have explored the theoretical foundations of NE algorithms such as their consistency for visualizing clustered data [15, 48, 2] and the existence of global minimizer of the loss function [4, 35]. However, there remains a lack of systematic evaluation and in-depth understanding about the key factors underlying the performance of NE algorithms in capturing smooth structures.

Dynamic modeling approaches based on biophysical principles and stochastic differential equations have also been developed to depict progression of cell-state transitions [89, 64, 79, 61, 77, 36, 75, 23]. While these methods have uncovered important biological insights on cell-fate dynamics, their efficacy relies heavily on the validity of the underlying assumptions about the dynamic system. A poor fit between the model and the observed data – for instance, when the model assumptions overlook a key factor – can significantly impair the accuracy of inferred dynamics and obscure meaningful biological signals. As such, it is crucial to incorporate a built-in model-checking procedure, to ensure the algorithm robustness and reliability. Furthermore, many of these algorithms rely on emerging data modalities to infer cell-state dynamics, such as RNA velocity [43, 79], cell lineage [77, 36, 75], experimental time points [64], and metabolic-labeling data [61]. While valuable in specific contexts, these approaches’ reliance on specific new data modalities limits their broader applicability, especially for analyzing the large volumes of existing single-cell data modalities, including scRNA-seq data, already generated from diverse biological systems. This underscores the need for general methods that can leverage the wealth of existing single-cell data and also are applicable to new data modalities to provide accurate insights into cell-state dynamics.

In this work, we introduce a novel machine learning approach to improve NE algorithms for reliably discovering low-dimensional smooth structures in single-cell data. Our approach is based on the PCS framework for veridical data science [85, 87, 84, 86], which employs core principles of Predictability, Computability, and Stability to provide formal guidelines for the entire data science life cycle, including method development, empirical evaluation and enhancement of trustworthiness in data analysis. Our contribution is two-fold. First, we conduct a systematic PCS-guided assessment of popular NE algorithms such as t-SNE, UMAP, PHATE, and a few others. Our evaluation integrates benchmark biological datasets with biological labels, numerical simulations, and a rigorous theoretical analysis to provide a comprehensive reality check (or “Predictability” assessment or Pred-check) for several popular NE algorithms. As a result, we uncover fundamental insights into their limitations in capturing low-dimensional smooth structures, including algorithmic artifacts and instability. Second, we propose a novel machine learning approach, NESS, which enhances <u>Neighbor Embedding</u> algorithms for <u>Smooth</u> structures using <u>Stability</u> measures. The proposed NESS approach leverages the intrinsic stochasticity of the NE algorithms, offered by the random initialization, to generate a series of low-dimensional embeddings and use the similarity between the embeddings’ local neighborhood structure to define stability measures for each data point. The NESS approach aims at improving diverse NE algorithms for modeling smooth structures in singlecell data, as well as providing biological insights on cell state dynamics. In particular, it may help identify stable and transitional cell states along developmental trajectories and quantify cellular transcription activity dynamics across different cell states.

To demonstrate its efficacy, we use NESS combined with t-SNE or UMAP to analyze six singlecell datasets (see Supplement Table S1 for each dataset and their abbreviations). These datasets cover diverse biological processes including hematopoiesis [59], human induced pluripotent stem cells (iPSC) differentiation [5], murine intestinal organoid development [6], dentate gyrus neurogenesis [31], spermatogenesis [29], and endoderm development in embryoid bodies [55]. They also encompass various sequencing technologies, such as scRNA-seq (10x Genomics Chromium), singlecell EU-seq, and single-cell RT-qPCR. Using biological labels from trusted external sources, we demonstrate that NESS (combined with t-SNE or UMAP) identifies both transitional and stable cell states across iPSC differentiation, organoid development, and embryoid body formation, predicts key genes involved in cell-state transitions, and quantifies cell-specific transcriptional activity in neurogenesis and spermatogenesis. Notably, our analysis resolves key neuronal subpopulations during embryoid formation and reveals novel biological insights into their respective cell states. Overall, our analysis suggests that NESS improves on NE algorithms to obtain more reliable characterizations of smooth biological structures in single-cell data.

## 2 Results

### PCS framework, overview of our approach, and NESS

The PCS framework for veridical data science [85, 87, 84, 86] is a principled data science workflow designed to enhance the reliability and interpretability of data-driven scientific discoveries. It is built upon three key principles: Predictability (P), Computability (C), and Stability (S). Specifically, the predictability principle ensures that models fit real-world data well, providing a comprehensive reality check using benchmark datasets to validate their usefulness and generalizability. The computability principle emphasizes efficient, reproducible and scalable computational implementations, and real-world-data-inspired simulations, ensuring coding robustness in practical applications. The stability principle assesses the sensitivity of results to reasonable data perturbations, model choices, and algorithmic randomness, helping to distinguish genuine findings from artifacts. The PCS framework integrates these principles throughout the entire data science pipeline, from problem formulation to method development, empirical evaluation, and interpretation. Our proposed approach, whose key idea and workflows are illustrated in Figure 1, is developed under the PCS framework, aiming to enhance the reliability of the analysis and interpretation of single-cell datasets with smooth structures from NE algorithms.

**Figure 1.**
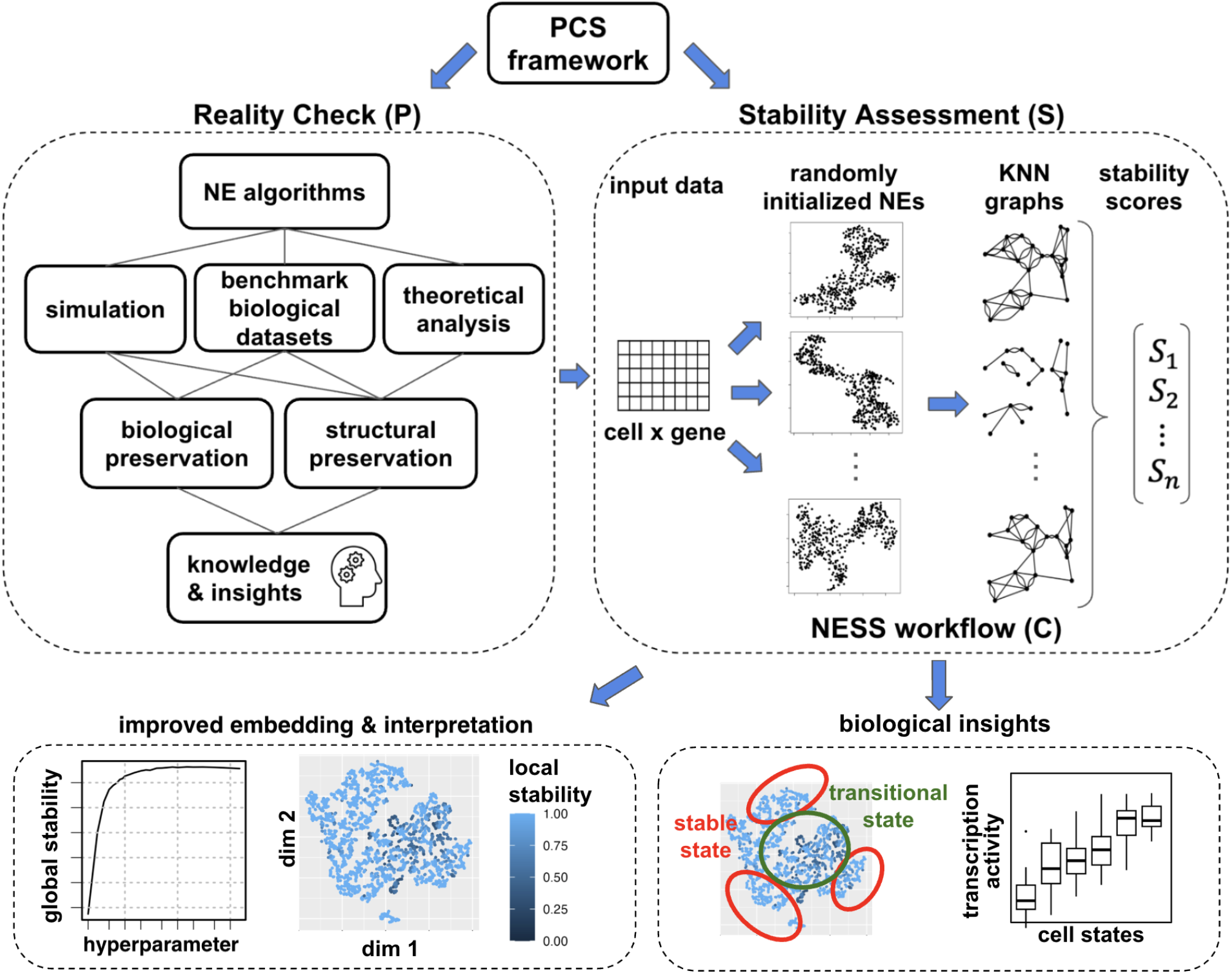
Overview and illustration of the proposed approach. Our approach is based on the PCS (Predictability, Computability, and Stability) framework for veridical data science. The “P” in PCS requires a comprehensive reality check for popular NE algorithms. Using numerical simulations, benchmark biological datasets, and theoretical analysis, we evaluate their performance and identify key limitations in capturing smooth structures and retaining biological signals. Guided by the “S” in PCS, we develop NESS, a novel algorithm for stability assessment and improvement of any NE algorithm on a given dataset without requiring ground-truth biological labels. NESS is computationally efficient (“C” in PCS). It leverages the random initialization of NE algorithms to generate multiple low-dimensional embeddings. From each embedding, it constructs a KNN graph among the data points (cells) and compares these graphs to compute a local stability score for each data point. In addition, NESS also generates a refined low-dimensional embedding that better reflects the latent smooth structure and provides an intuitive line chart with an automated workflow for hyperparameter selection. Building on these outputs, NESS aims to offer biological insights into cell progression dynamics, such as identifying stable and transitional cell states along developmental trajectories and quantifying transcriptional dynamics across cell states.

Following the PCS framework, we first perform a comprehensive reality check for popular NE algorithms under “P”, to evaluate their performance and identify key limitations in capturing smooth structures and retaining biological signals. We subject the low-dimensional embedding generated by each NE algorithm to several internal and external evaluation and validation checks. Internal validation uses benchmark single-cell datasets, numerical simulations, and/or rigorous theoretical analysis, to assess how well the embedding preserves local and global structure relative to the original high-dimensional data. External validation leverages biological labels (e.g., cell type or cell state annotations) associated with the benchmark datasets, available from trusted sources, to evaluate how well the NE algorithms retain biological signals. As detailed below, our systematic assessment reveals key insights into the limitations of the existing approaches and the main factors determining the performance of NE algorithms in characterizing smooth structures. These limitations include the artificial fragmentation of smooth structures within the embeddings and instability in the algorithm’s output with respect to random initializations.

To address these limitations, building on the insights from the internal and external checks, we propose a new algorithm, NESS, which provides stability assessment (“S” in PCS) of any NE algorithm with respect to a given dataset, aiming to enhance the performance of the NE algorithm in capturing the underlying smooth structure without requiring ground-truth biological labels. NESS exploits the inherent stochasticity of an NE algorithm, arising from random initialization of any iterative procedure, to introduce a point-wise stability measure for the final low-dimensional embeddings. Specifically, for a given input dataset and an NE algorithm, NESS first generates multiple low-dimensional embeddings of the dataset under different random initializations. From each embedding, we construct a *k*-nearest neighbor (KNN) graph among the data points (cells), and then compare the KNN graphs across all embeddings to assess the stability of each data point’s neighbors; see Methods for more details. As a result, NESS defines a local stability score for each data point, with a higher value indicating more consistent neighbors across multiple embeddings, providing a quantitative measure of the uncertainty in the obtained low-dimensional embeddings. Furthermore, NESS produces two additional outputs: a refined low-dimensional embedding that better reflects the underlying smooth structure, and an intuitive line chart with an automated workflow that guides hyperparameter selection. Building on these outputs, NESS aims to provide biological insights into cell progression dynamics, as supported from our analysis of six single-cell datasets. This includes using local stability scores to identify stable and transitional cell states along developmental trajectories together with the associated genes and pathways (Fig 4), and constructing a cell-specific transcription activity score to quantify the transcriptional dynamics across different cell states (Fig 5). Moreover, NESS is computationally efficient and scalable to large single-cell datasets with tens of thousands of cells (“C” in PCS). It has been implemented as an R package, available at our GitHub page https://github.com/Cathylixi/NESS, accompanied by PCS documentation and tutorials for broader community use.

### PCS-guided assessment reveals limitations of NE algorithms for smooth biological structures

Our PCS-guided assessment and validation of NE algorithms identify common limitations of popular NE algorithms, and provide empirical evidence and theoretical explanations of their unfavorable distortions of latent smooth structures. Moreover, our analysis offers key insights into the methodological strategies underlying NESS, which is to develop stability measures to improve NE algorithms.

#### Low graph connectivity is problematic for an NE algorithm to capture smooth structures

An important component in the existing NE algorithms is the construction of a neighborhood graph based on the input dataset. In each NE algorithm, the connectivity of the resulting graph is either explicitly or implicitly determined by a hyperparameter, hereafter universally referred to as the “graph connectivity parameter” (GCP), that controls the resolution of the neighborhood definition. For example, in UMAP, densMAP, and PHATE (hereafter referred to as PhateR), the GCP is defined as the number of neighbors *k* in the *k*-nearest neighbor graph based on the input data [54, 55, 56]; in t-SNE, it is defined as the “perplexity,” which determines edge weight assignment through entropy measures [71]. In general, low GCP values produce sparser graphs, reducing computational cost, whereas high GCP values lead to denser graphs with increased computational demands. In practice, smaller GCPs are often preferred for efficiency, yet there is no principled approach for selecting GCPs that best preserve smooth structures.

To evaluate NE algorithms relative to their GCP hyperparameter settings, we carry out a systematic reality check of their performance on both synthetic data and real-world single-cell datasets, each representing a distinct developmental process, including tissue-specific lineage trajectories (Mouse Hema), pluripotent stem cell differentiation (iPSC), and organoid development (Murine Intestinal). Every single-cell dataset contains appropriate biological labels such as cell types or time points (indicative of cell state). We assessed four widely used NE algorithms, including t-SNE, UMAP, PhateR, and densMAP, using multiple evaluation metrics. To quantify structure preservation, we measure the neighborhood concordance and the Pearson correlation of pairwise distances between the low-dimensional embedding and input data (Methods). To assess the preservation of biological signals, we use the Silhouette index, neighbor purity index, and local Simpson index, in each case comparing the embedding coordinates with biological labels (Methods). Our results highlight the critical role of GCP selection in ensuring reliable embeddings (Figures 2, S1–S3). Specifically, we find that for all NE algorithms evaluated in this study, regardless their initialization schemes, under-specifying GCPs tend to introduce substantial distortions, causing artificial fragmentation and disruptions in the latent smooth structures (Figures 2a, S1–S3). For example, for our synthetic data (Simulated Curve) randomly sampled from a one-dimensional curve representing an ideal, noiseless progression trajectory, t-SNE embeddings with low GCP values fragmented the continuous curve into multiple disconnected segments; in the Mouse Hema dataset, low-GCP t-SNE embeddings fragmented the two differentiation branches into many small, disconnected clusters, obscuring the true trajectory structure. See Supplementary Figures S1–S3 for additional results under different initialization schemes and dataset-method combinations. Conversely, except for PhateR, which exhibits a slight decline in performance across various metrics under very large GCPs, most NE algorithms display robust performance with higher GCPs, and are relatively insensitive to GCP variations in that range (Figure 2b). The slight performance decay of PhateR under large GCPs is likely due to a mild over-smoothing effect, causing data points to concentrate along certain learned trajectories while still preserving the underlying latent structure (Supplement Figure S4). However, as expected, the benefits of higher GCPs come at the expense of increased computational time (Supplement Figure S5a). As such, an important benefit of NESS, as detailed below, is to address such limitations and provide an automated selection of GCPs that balances computational cost and embedding faithfulness.

**Figure 2.**
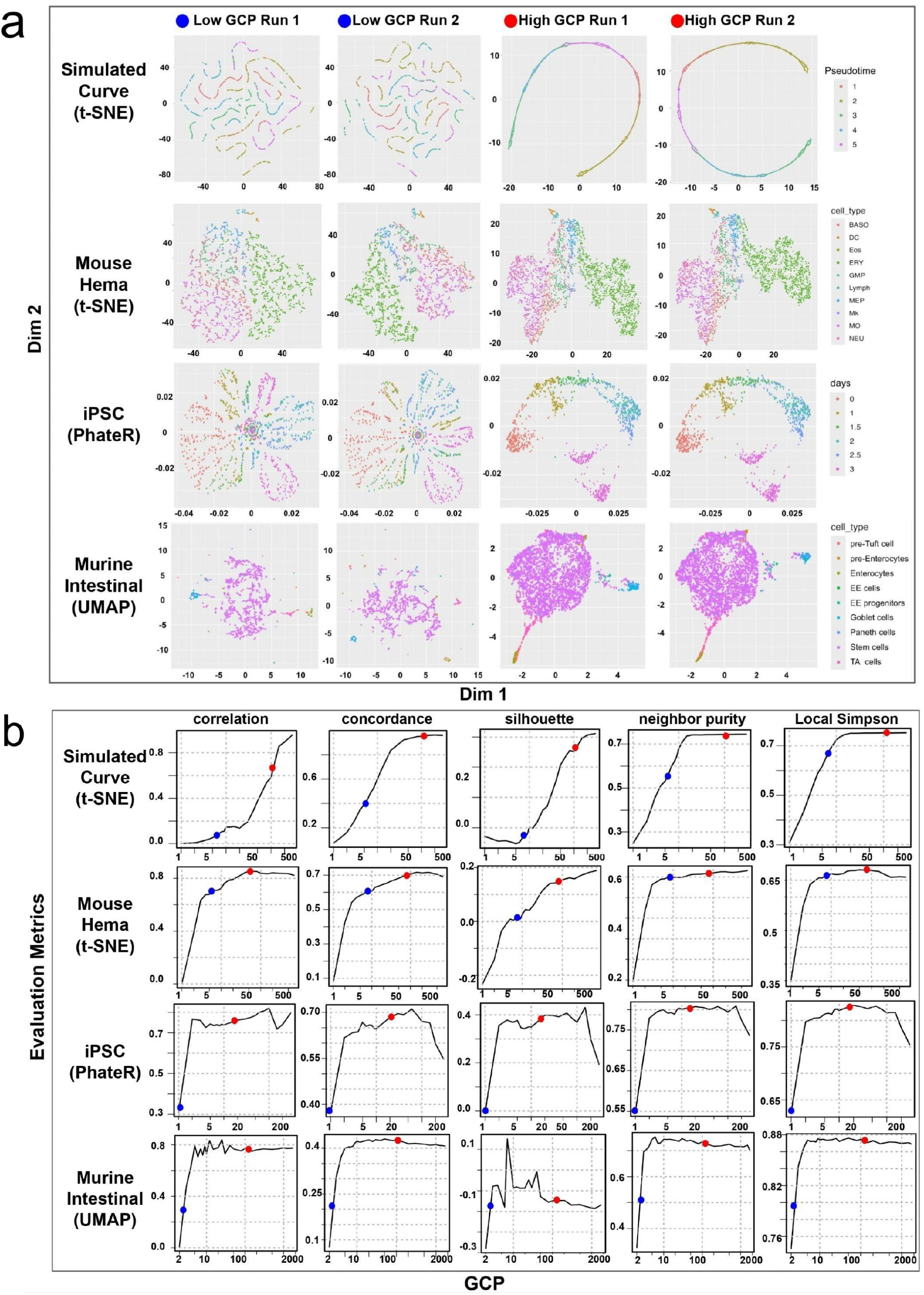
PCS-guided assessment identifies artifacts in NE algorithms. (a) Examples of low-dimensional embeddings for synthetic (first row) and benchmark single-cell datasets (last three rows) generated by popular NE algorithms, including t-SNE, UMAP, and PhateR, with cells colored by their labels. The left two columns correspond to two instances of RIs under relative low GCPs, whereas the right two columns show embeddings under relatively high GCP values. The specific low and high GCP values are indicated in (b). See Methods for the definition of GCP in each NE algorithm. All embeddings are generated using random initialization (RI). See Supplementary Figures S1–S3 for more examples and similar results under alternative initialization schemes, and Supplementary Figure S6 for the ground truth cell differentiation hierarchies. (b) Assessment of NE algorithm performance based on multiple metrics and the effect of GCP tuning. Line charts depict the relationship between GCP (x-axis, log scale) and five internal and external evaluation metrics (y-axis): Correlation, Concordance, Silhouette Score, Neighbor Purity, and Local Simpson Index, where higher values indicate more reliable embeddings. The red dots and blue dots on the line charts indicate the respective high and low GCPs in (a). Our results suggest underspecifying GCP tends to introduce artificial fragmentation and disruption of the latent smooth structure, while increasing instability in the final embeddings. In contrast, embeddings under higher GCPs reliably reveal cell differentiation hierarchies.

#### Theoretical analysis of t-SNE connects low graph connectivity to fragmented embeddings

In addition to the empirical assessments above, we also establish a mathematical connection between low GCP values and artifacts in low-dimensional embeddings produced by NE algorithms. In particular, through rigorous analysis of t-SNE, we find that for synthetic data generated from a low-dimensional manifold model, an under-specified GCP inevitably leads to a pathological deformation of the optimization landscape, causing severe distortions in the final embedding (Theorem 4.1 in Methods). Moreover, our analysis highlights that these artifacts arise primarily from discontinuities or “gaps” in the low-dimensional representation, explaining the fragmented patterns observed in various examples (Figures 2a and S1, left two columns, and Figure 3a middle column). Despite that our theoretical analysis only focuses on t-SNE, the underlying similarity between the loss functions of t-SNE and other NE algorithms [12, 38, 21] suggests that our findings likely extend to a broader class of NE methods.

**Figure 3.**
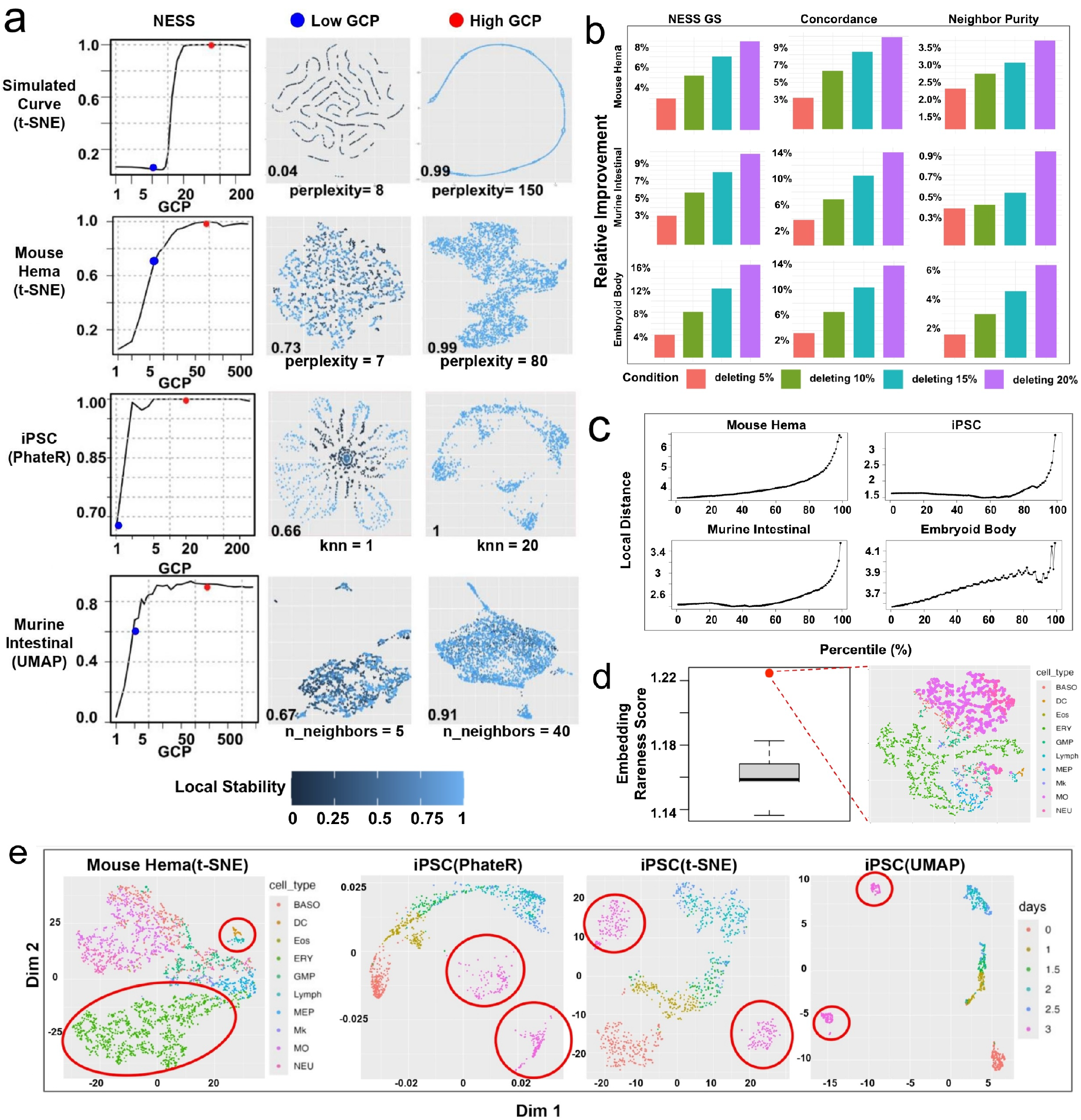
NESS enhances representation and uncertainty quantification of smooth structures. (a) Left column: line charts showing the relationship between GCP tuning and the NESS global stability (GS) score for various NE algorithms, evaluated on simulated and real single-cell datasets. Middle and right columns: low-dimensional embeddings corresponding to two GCP values, a lower GCP value (“Low GCP”) and a NESS recommended GCP value (“High GCP”), indicated by blue and red dots respectively in the line charts. The GCP and its value for each NE algorithm are indicated below the embeddings. Data points are colored by NESS local stability scores, with the corresponding GS scores shown in the bottom-right corner of each plot. Higher GCP values generally improve stability and structure preservation. Supplementary Figures S7 and S8 illustrate how these stability measures can also guide the selection of NE methods and other hyperparameters. (b) Barplots showing the relative improvement in NESS GS score, concordance, and neighbor purity across different datasets as increasing percentages (5%, 10%, 15%, 20%) of low-stability cells are removed. All GS scores are evaluated under the respective NESS recommended GCP values in (a). The iPSC dataset is excluded since its global stability reaches 1. (c) Cells with low NESS local stability scores tend to have greater distances from their neighboring cells. Plots show the average distance between each cell and its 30-nearest neighbors, among cells whose local stability score falls in the *p*% upper percentile of the cells, all obtained under NESS recommended GCPs. Higher percentiles correspond to cells with lower NESS local stability and increased local distances to their neighbors, indicating their tendency to reside in low-density regions. (d) Left: Boxplot of embedding rareness scores for 50 t-SNE embeddings of the Mouse Hema dataset under RI and the NESS recommended GCP in (a), with the largest score marked in red. Right: t-SNE embedding corresponding to the largest embedding rareness score, with cells color-coded by cell type. Structural distortions in monocytes (purple color) and neutrophils (red color) are highlighted, where originally neighboring cell populations are separated into distinct regions. (e) Examples of low-dimensional embeddings obtained by NE algorithms under their default GCPs. First panel: t-SNE under default GCP (GCP=30) applied to the Mouse Hema dataset led to artificial fragmentation of erythrocytes (ERY) and separated dendritic cells (DC) from the main developmental trajectory, as highlighted in red circles. Last three panels: default applications of PhateR (default GCP=5), t-SNE (default GCP=30), and UMAP (default GCP=15) to the iPSC dataset fail to capture the lineage relationship (Supplementary Figure S6b) between the primitive streak population at day 2-2.5 and the differentiated mesodermal and endodermal cells at day 3, as highlighted in red circles. These artifacts can be overcome by NESS, as shown in the second and third rows of Figures 2a and 3a under “High GCP”, which is the NESS recommended GCP.

**Figure 4.**
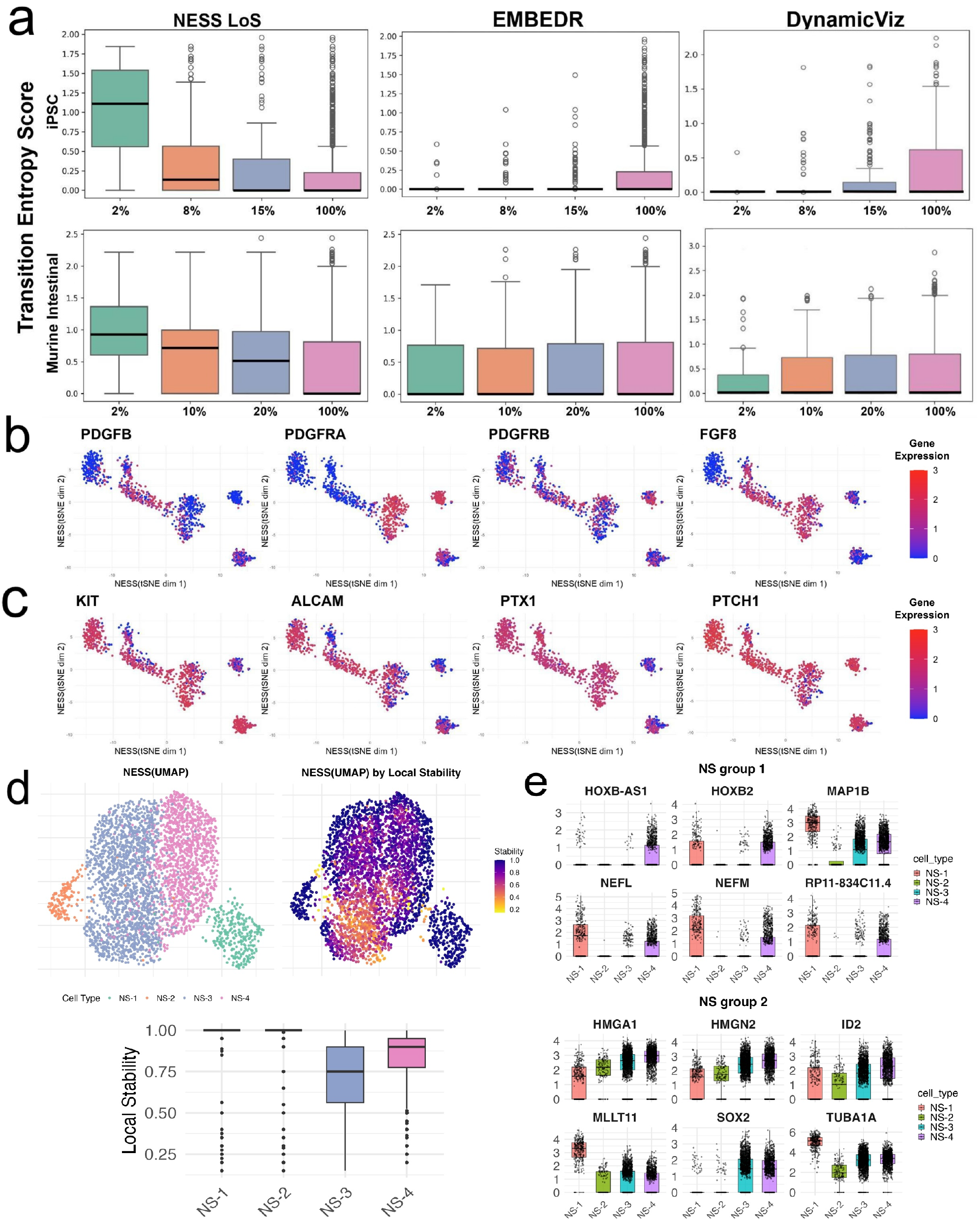
NESS identifies transitional and stable cell states in diverse biological developments. (a) Boxplots of MuTrans’ transition entropy scores for cells whose NESS(t-SNE) local stability (LoS) score, and the pointwise stability measures from EMBEDR, and DynamicViz (see Methods) are at the bottom 2%, 8%, 15% and 100% percentile. Higher entropy score indicates greater likelihood of being in a transition cell state. Top row: iPSC dataset. Bottom row: Murine Intestinal dataset. Cells with low NESS local stability scores tend to have higher transition entropy score. These results suggest the advantage of NESS local stability score in consistently identifying transitional cell states compared with EMBEDR and DynamicViz. (b) NESS enhanced t-SNE embeddings of the iPSC dataset, colored by the expression level of *PDGFB, PDGFRA, PDGFRB*, and *FGF8*, whose expression is negatively correlated with NESS local stability. (c) NESS-assisted t-SNE embeddings of the iPSC dataset, colored by the expression level of *KIT, ALCAM, PTX1*, and *PTCH1*, whose expression is positively correlated with NESS local stability. (d) NESS-assisted UMAP visualization of neuronal subtypes NS-1 to NS-4 from the Embryoid Body dataset The top left panel shows cell types, while the top right panel displays NESS local stability scores. Notably, NS-3 and NS-4 cells exhibit lower local stability. (e) Boxplots of gene expression in NS-1 to NS-4 cells. Genes are grouped into two categories based on expression patterns: Group 1 includes genes for which NS-4 expression resembles NS-1, while Group 2 includes genes for which NS-4 expression resembles NS-3. These results suggest that NS-4 and NS-3 represent progenitor-like or intermediate cell types, while NS-1 and NS-2 exhibit more stable, differentiated identities, consistent with their local stability scores, respectively.

**Figure 5.**
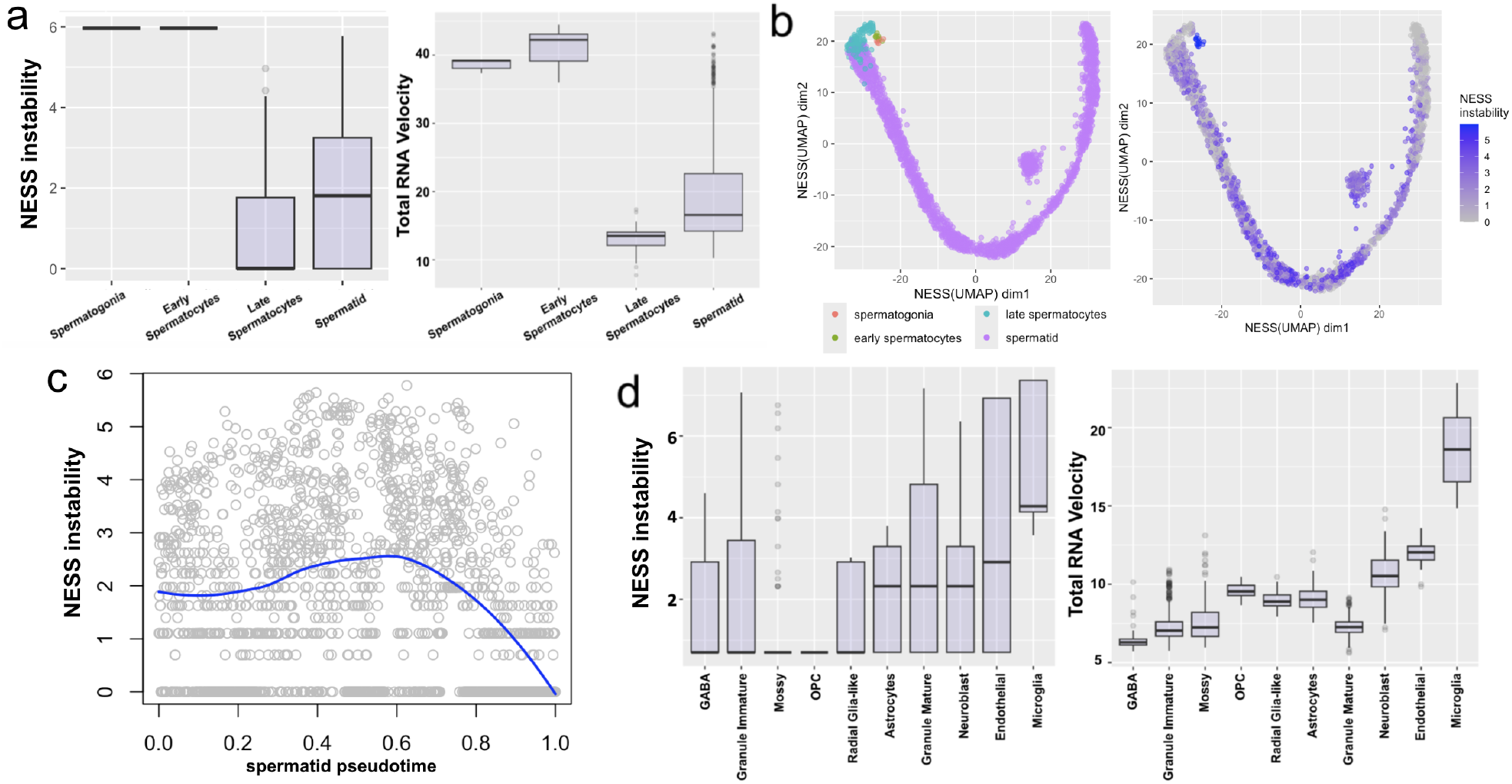
NESS reveals transcriptional dynamics during spermatogenesis and neurogenesis. (a) Comparison of pointwise NESS instability score, defined as 1/(NESS local stability score), with total RNA velocity for the Spermatogenesis dataset. Each boxplot contains the cells within a specific cell type. The total RNA velocity is defined as the *L*_2_-norm of the RNA velocity vector across the top 2000 most variable genes, independently obtained by scVelo, which incorporate additional information from RNA splicing dynamics. The plots demonstrate remarkable similarity between the patterns of NESS-TAG scores and the total RNA velocity across different cell tyles. (b) NESS-assisted UMAP visualization of the Spermatogenesis dataset, colored by cell types (left) and NESS instability score (right). (c) NESS instability score of the spermatid cells ordered by pseudotime along the cell progression trajectory in (b). The blue line indicates the loess fitted regression curve. (d) Comparison of NESS instability score with total RNA velocity for the Dentate gyrus neurogenesis dataset. The boxplots are ordered by the median of NESS instability within each cell type. The results indicate notable concordance between the patterns of the two measures across cell types.

#### Random initialization provides a natural and computationally efficient form of perturbation for stability analysis

Another crucial factor of the NE algorithms is the initialization scheme [38]. As iterative algorithms, NE methods commonly use random initialization (RI), PCA initialization, Laplacian initialization, and so on. Among them, RI, which generates the initial embedding coordinates by randomly sampling within a small region around the origin of the low-dimensional space [71], has often been criticized for its lack of interpretation, and potential to misrepresent global structures [15, 38]. However, inspired by the PCS framework, we view the RI as a natural tool for creating algorithmic perturbations, and to generate stability assessment for the NE algorithms. Our findings reveal that low GCP values, combined with RI, lead to highly variable distortions across different runs (Figure 2a and Supplement Figure S1). For example, in the synthetic data (Simulated Curve), the low-GCP t-SNE embeddings contain arbitrary fragmentation and an arbitrary layout of the underlying curve; for the iPSC dataset, both low-GCP PhateR embeddings failed to capture the latent differentiation trajectory, yet displayed flower-shaped patterns, with notable structural discrepancies between the resulting embeddings. Similar fragmentation and instability are observed in the Mouse Hema dataset visualized with t-SNE and the Murine Intestinal dataset visualized with UMAP. In contrast, NE embeddings under relatively higher GCPs exhibit greater structural stability under RI. These findings suggest that careful selection of GCP not only enhances embedding quality but also improves the algorithm’s robustness to initialization variability. This key insight underpins the main idea behind the design of NESS, which is to develop stability measures to achieve more reliable performance.

### Proposed NESS enhances NE representation and stability assessment of smooth biological structures

To mitigate potential artifacts of NE algorithms in single-cell analysis, where ground-truth biological labels are typically unavailable, we propose the NESS approach to enhance the representation and stability assessment of smooth biological structures based on the stability principle. We demonstrate that NESS improves the performance of diverse NE algorithm in characterizing smooth biological structures in four key aspects.

#### NESS global stability score optimizes embedding of smooth structures and guides hyperparameter selection

For a given dataset and an NE algorithm, NESS first defines a local stability score for each cell and uses its average across all cells to define a global stability (GS) score of this NE algorithm (Methods). It then generates a line chart where the x-axis is the GCP value and the y-axis is the global stability scores of the NE algorithm. It facilitates improved and automated selection of suitable GCPs in an unsupervised manner (i.e., without biological labels). From the line chart, users can select a GCP associated with a higher GS value while avoiding unnecessary computational overhead that only marginally improves embedding quality (Fig. 3a, left column). To further enhance computational efficiency and practicality, we develop a workflow (Methods) that automatically selects a suitable GCP for downstream analysis. It essentially recommends the smallest GCP that achieves a high GS score or shows no significant improvement over a slightly smaller GCP. We find that NESS effectively mitigates structural distortions across various NE algorithms (Fig 3a, middle column), leading to improved visualizations of smooth biological structures in different contexts (Fig 3a, right column). For example, NESS successfully generates informative embeddings that reveal key differentiation pathways, including the transition of mouse HSC cells into lymphoid and myeloid lineages (Figures 2a and 3a second row, and Supplement Figure S6a), the differentiation of iPSC cells into endodermal and mesodermal lineages (Figures 2a and 3a third row, and Supplement Figure S6b), and the progression of murine intestinal stem cells into secretory cells and enterocytes (Figures 2a and 3a fourth row, and Supplement Figure S6c). Similarly, the GS metric can be applied to compare and optimize other hyperparameters, including categorical ones such as the choice of distance metric in a given NE algorithm (Supplementary Figure S7). Additionally, it can aid in selecting among different NE algorithms, helping to identify the most suitable method for visualizing a specific dataset (Supplementary Figure S8).

#### NESS local stability score quantifies pointwise embedding reliability

For any NE algorithm with NESS recommended hyperparameters, the NESS pointwise local stability score quantifies the algorithm’s technical stability on a given dataset. These scores help distinguish more stable regions of the embedding from those that are less stable, providing insight into where the algorithm consistently recovers structure. Importantly, regions with low stability scores may also correspond to areas of higher biological uncertainty, where the underlying structure is more ambiguous or less well-defined. By highlighting both algorithmic variability and potential ambiguity in the data, these measures support more informed interpretation of local structures in the embedding. For example, in the Murine Intestinal dataset, NESS(UMAP) under the recommended GCP value (GCP=40, or “High GCP” in Figure 3a), we identified a group of Enteroendocrine Progenitor (EE progenitors) cells whose low-dimensional embeddings exhibit greater local instability compared to other cells within the same embedding (Figure 3a, bottom right panel; Supplement Figure S9), indicating higher variability or biological uncertainty as compared with other cells. Additionally, in the Mouse Hema, Murine Intestinal, and Embryoid Body datasets, we find that removing the top 5–20% most unstable cells, identified by low NESS(t-SNE/UMAP) local stability scores (see Methods), consistently improves embedding quality. These unstable cells likely contain higher levels of noise or subtle biological signals not aligned with the primary trajectory. Their removal thus enhances both the preservation of smooth structural patterns (measured by Concordance) and the biological signal (measured by Neighbor Purity) in the remaining embeddings, resulting in improved overall stability (Figure 3b). The stratification of cells according to their local stability measures can thus refine the identification of global smooth structures. Moreover, across various biological datasets and different NE algorithms, we observe a strong association between NESS(t-SNE/UMAP) local stability and local cell density, where the cells with low stability tend to have greater distances from their nearest neighbors than expected (Figure 3d, Methods). This suggests that highly unstable cells often reside in low-density regions of the feature space. If we assume continuity of the cell state changes, such low-density regions would therefore consist of cells undergoing fast state transitions. This connection allows us to interpret NESS local stability scores not only as a measure of the technical stability of NE algorithms but also as an indicator of the biological stability of cell states, providing valuable biological insights, as discussed below.

#### NESS embedding rareness score avoids technical artifacts from RI

For any NE algorithm with NESS recommended hyperparameters, in order to choose a suitable low-dimensional embedding among the multiple (e.g., 50) embeddings obtained under different RIs, NESS also generates an “embedding rareness” score for each embedding, quantifying how similar each embedding is to the ensemble of embeddings generated from multiple RIs. It has been observed that even with a well-chosen GCP, a poor embedding may occasionally arise by mere chance from RI [15]. For example, when using the same recommended GCP as in the right panel of Figure 3a, a low-dimensional embedding produced by t-SNE can still exhibit substantial distortions (Figure 3c, right). In this case, both the monocyte and neutrophil populations–originally close neighbors in the high-dimensional feature space–are fragmented into disconnected regions in the embedding. The rareness score can thus help avoid such pathological cases caused by RI, ensuring selecting more reliable embeddings for better visualization (Figure 3c, left).

#### NESS-assisted NE algorithms improves substantively standard NE algorithms under default GCPs

Lastly, we observe that while conventional application of NE algorithms under their respective default hyperparameters appeared to effectively capture the underlying biological structure in some datasets, such as Murine Intestinal dataset (Supplement Figure S11), it may introduce significant artifacts in other datasets. For example, applying t-SNE with its default GCP (`perplexity`=30 in R) to the Mouse Hema dataset led to artificial fragmentation of erythrocytes (ERY) into small clusters, and separated dendritic cells (DC) from the main developmental trajectory (Figure 3e, first panel, artifacts highlighted in red circles). Similarly, default applications of PhateR (default GCP `knn`=5 in R), t-SNE (default GCP `perplexity`=30 in R), and UMAP (default GCP `n neighbors`=15 in R) to the iPSC dataset generated low-dimensional embeddings that fail to capture the lineage relationship (Supplementary Figure S6b) between the primitive streak population at day 2-2.5, and the downstream differentiated endodermal and mesodermal cells at day 3 (Figure 3e, last three panels, artifacts highlighted in red circles). In contrast, NESS-assisted embeddings corrected these distortions–for example, in the NESS-assisted t-SNE embeddings of Mouse Hema dataset (the second rows of Figures 2a and 3a under “High GCP”, which is the NESS recommended GCP) and in the NESS-assisted PhateR embeddings of iPSC dataset (the third rows of Figures 2a and 3a under “High GCP”). These results highlight the benefits and improvements from integrating NESS into routine single-cell analysis pipelines that utilize NE algorithms.

#### NESS identifies transitional and stable cell states in diverse biological developments

We show that NESS combined with NE algorithms such as t-SNE or UMAP under the NESS recommended GCP identifies transitional and stable cell populations through its local stability scores, and provides biological insights into cellular differentiation and lineage progression.

#### Validating NESS local stability score using cell transition entropy and cell-state annotations

To validate our method, we use the transition entropy score computed by MuTrans [89], an algorithm that directly models cell-fate transition dynamics based on biophysical principles. MuTrans computes a transition entropy score for each cell, quantifying its likelihood of being in a transition state, with higher values indicating stronger evidence of intermediate cell states with mixed identities. We focus on the iPSC and Murine Intestinal datasets, on which MuTrans’ performance was certified in the original publication [89]. We validate NESS’s performance by comparing the local stability score from NESS(t-SNE) with MuTrans’ transition entropy scores. As another validation, we evaluate the local stability score from NESS(UMAP) on the Embryoid Body dataset where the detailed cell state annotations are available [55]. To benchmark the performance and demonstrate the advantages of NESS, we compare its results with existing embedding assessment algorithms, including EMBEDR, DynamicViz, and scDEED.

We observe a strong association between the NESS local stability (LoS) score and the transition entropy score. Specifically, in both iPSC and Murine Intestinal datasets, cells in the bottom percentile with lower local stability scores tend to have higher transition entropy scores, suggesting that the cells demonstrating greater variability in the NE embeddings are more likely in transitional states. In contrast, alternative embedding assessment methods, such as EMBEDR and DynamicViz (note that scDEED does not produce similar continuous assessment scores), do not exhibit similar associations (Figure 4a).

To further validate NESS, we apply it to the Embryoid Body dataset, comparing the NESS(UMAP) local stability score across a spectrum of cell types and cell states, from pluripotent stem cells (ESCs) to lineage-committed progenitors (Supplement Figures S6d and S10b). The annotated cell states and the lineage relationships among them were previously characterized and experimentally validated in [55]. Consistent with expected developmental hierarchies, intermediate cell states (e.g., NE-1/NS-5 and EN-1) exhibit lower local stability scores, while self-renewing ESCs and terminally differentiated populations (e.g., CPs, EPs, SMPs) display higher stability scores (Supplement Figures S10a). As a comparison, alternative methods such as EMBEDR and scDEED do not appear to distinguish transitional and stable cell states; while DynamicViz is able to similarly identify ESC, EPs and SMPs as stable cell states, it does not capture the relative stability of CPs (Supplement Figure S10a).

#### NESS helps identify key genes associated with iPSC differentiation

To demonstrate the implication of the NESS-identified transitional and stable cell states in downstream analysis, we evaluated associations between gene expression and the NESS(t-SNE) local stability across all cells in the iPSC dataset. Our analysis detected 48 key genes whose expressions are significantly associated with NESS local stability score (see Methods, Table S2). Among them, *PDGFB* (adjusted p-value = 3.17 × 10^−3^), *PDGFRA* (adjusted p-value = 1.4 × 10^−4^), *PDGFRB* (adjusted p-value = 3.07 × 10^−3^) and *FGF8* (adjusted p-value = 3.97 × 10^−9^) were top genes negatively correlated with NESS local stability. As expected, these genes display relatively higher expression in the less stable, transitional cell states (Figure 4b and S6b), highlighting both the transition from the initial epiblast to the primitive streak population (as in *PDGFB* and *PDGHRB*), and the subsequent differentiation of the primitive streak into endodermal and mesodermal cell types (as in *PDGFRA* and *FGF8*) [5]. In contrast, *KIT* (adjusted p-value = 2.08 × 10^−2^), *ALCAM* (adjusted p-value = 3.23 × 10^−4^), *PTX1* (adjusted p-value = 6.2 × 10^−4^) and *PCH1* (adjusted p-value = 3.45 × 10^−3^), among others, were positively correlated with NESS local stability score, showing higher expression in the more stable cell states (Figure 4c), which mainly consist the complementary, non-transitional populations. In particular, among genes whose expression is localized in transitional cell states, *PDGFB, PDGFRA* and *PDGFRB* are associated with the Platelet-Derived Growth Factor (PDGF) signaling pathway, which is essential for mesodermal differentiation, influencing vascular development, mesenchymal stem cell formation, and cardiac lineage commitment [11, 1, 30]; *FGF8* is a key regulator of neurodevelopment, driving the differentiation of iPSCs into neural progenitors\ and contributing to midbrain formation [19, 70, 57]. Among genes whose expression is localized in more stable iPSC cell states, *KIT* (c-KIT) is a cytokine receptor expressed on the surface of hematopoietic stem cells as well as other cell types, which plays a role in cell survival, proliferation, and differentiation [44]; activated leukocyte cell adhesion molecule (*ALCAM*) is known as a marker of fetal mouse and human iPSC-derived hepatic stellate cells [3, 40]. Moreover, the enrichment analysis of the identified 48 genes highlighted important pathways such as MAPK signaling (p-value = 3.4 × 10^−10^), notch signaling (p-value = 4.9 × 10^−7^), cytokine-cytokine receptor interaction (p-value=7.1×10^−4^), and Wnt signaling (p-value = 3.8×10^−3^). These are known to be key pathways in regulating iPSC proliferation, differentiation and reprogramming [25, 33, 73, 22]. These results demonstrate NESS’s ability to uncover biologically relevant transcriptional programs governing cell-state transitions.

#### NESS resolves distinct neuronal subpopulations during embryoid formation

Finally, we show that the NESS(UMAP) local stability score helps to resolve distinct neuronal subpopulations during embryoid formation and provides new insights into their cell states. In the Embryoid Body dataset, prior analyses identified four neuronal subtypes (NS-1 to NS-4) but were unable to resolve the finer structure of their respective cell states or the relationships among these subpopulations [55]. In our analysis, neuronal subtypes NS-3 and NS-4 display lower local stability scores compared to NS-1 and NS-2, suggesting the presence of subpopulations within NS-3 and NS-4 that demonstrate transcriptional mixing between them (Figure 4d). To further investigate the molecular basis of this instability, we performed differential gene expression analysis between NS-1/2 and NS-3/4, identifying genes with adjusted p-values less than 0.001. This analysis revealed two distinct gene groups (Figure 4e). In the first group, genes show higher expression in NS-1, moderate expression in NS-4, and minimal or no expression in NS-2 and NS-3; for example, we have *NEFM* (Neurofilament Medium Chain), which encodes an intermediate filament essential for maintaining neuronal caliber, axonal transport, and structural integrity of neurons [88]. In the second group, genes exhibit similar expression patterns between NS-3 and NS-4 but are distinctly expressed compared to NS-1 and NS-2. An example is *SOX2*, a transcription factor essential for maintaining neural progenitor identity and regulating early neural differentiation [78], which is highly expressed in NS-4.

This suggests that NS-4 may correspond to a progenitor-like population. NS-3 displays intermediate expression patterns, consistent with a transitional cell state, while NS-1 and NS-2 express neuronal maturation markers such as *NEFM*, indicating more differentiated identities. Together, these observations support a model in which NS-4 and NS-3 represent transcriptionally unstable or transitional populations, aligning with their lower local stability scores.

### NESS reveals transcriptional dynamics during spermatogenesis and neurogenesis

Building upon the above connection between the NESS local stability score and cellular transitivity, with the assumption that cells undergoing cell-state transitions likely exhibit a higher rate of change in the transcriptional activity, we show that NESS combined with t-SNE or UMAP can also serve as a proxy for inferring transcriptional dynamics using only scRNA-seq data. To illustrate this utility, we analyze two additional scRNA-seq datasets, one on the sermatogenesis, and the other on the neurogenesis, with NESS(UMAP). As a validation step, we compare the NESS local stability score with cell-specific RNA velocity estimates independently obtained using scVelo [9], which incorporate additional information from RNA splicing kinetics to quantify transcriptional activity. In particular, for each cell, we define total RNA velocity as the *L*_2_-norm of the estimated RNA velocity vector across the top 2,000 most variable genes, quantifying the overall rate of change in transcription activity.

For the Spermatogenesis dataset, our analysis reveals a remarkable similarity between the patterns of pointwise NESS instability score, defined as 1/(NESS local stability score), and the total RNA velocity. Among the four major stages of spermatogenesis, cells in the spermatogonia and early spermatocyte stages exhibit greater NESS local instability and higher total RNA velocity, while cells in the late spermatocyte and spermatid stages display lower and moderate values, respectively (Figure 5a, b). This pattern aligns with current biological understanding of germ cell development. Specifically, spermatogonia and early spermatocytes undergo rapid mitotic divisions and initiate meiosis, likely contributing to elevated transcriptional activity [27]. In contrast, as cells progress to late spermatocytes and spermatids, global transcription tend to decline and stabilize, as post-meiotic cells undergo extensive morphological changes, driven largely by pre-synthesized transcripts and post-transcriptional regulation [13]. Notably, within the spermatid subpopulation, we observe a gradual rise and subsequent decline in NESS instability score along the progression trajectory (Figure 5b, c). This dynamic likely reflects the transition from active gene expression in early spermatogenesis to transcriptional quiescence in later stages, facilitating efficient germ cell maturation through post-transcriptional mechanisms.

For the Dentate gyrus neurogenesis dataset, we also observe notable concordance between the patterns of NESS(UMAP) instability score and the total RNA velocity across different cell states (Figure 5d). The greater variability in the NESS instability score within each cell type compared with the total RNA velocity is likely due to the higher noise level in scRNA-seq data when inferring transcription dynamics, relative to the spliced RNA data used to estimate RNA velocity. In particular, our analysis highlights the relatively higher transcriptional activity of microglia and endothelial cells, while GABAergic neurons and immature granule cells exhibit lower transcriptional activity. These patterns align with known cellular functions and developmental dynamics of neurogenesis. Specifically, microglia and endothelial cells play crucial roles in immune response, neuroinflammation, and vascular remodeling, necessitating continuous gene expression and rapid transcriptional adaptation [53, 20]. In contrast, immature granule cells are known to develop very slowly [90], allowing for greater plasticity and adaptability during their maturation phase [58]. Similarly, GABAergic neurons, which are largely post-mitotic, experience transcriptional stabilization following differentiation [46]. These findings underscore the biological relevance of NESS in capturing transcriptional dynamics across different cell states in neurogenesis.

## 3 Discussion

Overall, we develop a unified approach NESS for assessing and improving NE algorithms based on PCS principles, enabling more reliable visualization and interpretation of smooth structures in single-cell data while offering valuable biological insights into cell development. A key strength of the NESS local stability score lies in its design principle—leveraging random initialization to introduce algorithmic perturbations, allowing a systematic evaluation of embedding stability. Compared with existing embedding evaluation algorithms, NESS shows three key advantages. First, the NESS evaluation is not restricted to any particular instance of embedding. In contrast, a baseline method scDEED assesses neighborhood structures by comparing a single realization of a (potentially random) low-dimensional embedding with the original dataset. Consequently, the inferential insights from scDEED are limited to that particular embedding instance and may not readily generate additional insights into the input dataset, such as local density variations and cell transition stability. Second, NESS’s algorithmic perturbation strategy is more effective at handling rare cell types and computationally more efficient than sample-perturbation-based methods. For example, approaches like DynamicViz and EMBEDR generate uncertainty estimates by resampling the entire input data. While this enables assessment of embedding variability, such methods tend to emphasize well-represented cell states with sufficient sample sizes, often underrepresenting rare populations due to low sampling rate. Finally, NESS is broadly applicable and can be integrated with any NE algorithm. In contrast, techniques like LOO require detailed knowledge of specific embedding algorithms, such as their objective functions and explicit Hessian expressions, which restricts their utility to a narrow subset of methods such as t-SNE and UMAP.

Compared with dynamic modeling algorithms, NESS also has the following conceptual and practical advantages. First, unlike methods that rely on strict mathematical modeling of cell-state transitions, NESS is inherently more flexible with respect to data-generating mechanisms and latent biological processes. Second, most existing dynamic modeling algorithms, such as scVelo, Velocyto, and MuTrans, lack model-fitting assessment options. In contrast, NESS provides multiple assessment measures to ensure the reliability of analysis results. Third, due to its simpler algorithmic design, NESS requires lower computational costs and exhibits superior scalability for large datasets with over 10^4^ cells. Across multiple single-cell datasets of varying sizes, our analysis yields that NESS significantly outperforms MuTrans in computational efficiency, requiring no more than one-third of MuTrans’s computing time for any datasets with over 3,000 cells (Supplement Figure S5b).

NESS has some limitations that we plan to address in future work. First, one assumption behind NESS is that the underlying smooth structure of interest should be intrinsically low-dimensional, so that the low-dimensional embeddings do not face the fundamental impossibility of embedding a higher-dimensional manifold into a lower-dimensional space. This assumption may be violated in complex biological processes where cell development cannot be adequately represented by a lowdimensional cell-state manifold. To address this challenge, we aim to develop methods for estimating the intrinsic dimensionality of high-dimensional datasets, providing a principled way to assess the suitability of NESS in a given context. Second, NESS relies on existing NE algorithms to generate low-dimensional embeddings and is thus inherently constrained by their individual limitations, even though it can select the most suitable algorithm among them (Supplement Figure S8). A promising next step is to develop an ensemble approach [52] that, after reality-check through “P” in PCS, combines the strengths of different Pred-checked NE algorithms as well as other embedding algorithms, such as Laplacian eigenmap [7] and diffusion map [42], to improve the overall quality of the low-dimensional embeddings.

## 4 Methods

### 4.1 PCS-Guided Assessment and Validation of NE Algorithms

Our PCS-guided reality check of NE algorithms involves two parts. On the one hand, we validate the preservation of biological information in the low-dimensional embedding using known biological labels of the cells, such as cell type and cell state annotations, associated with the benchmark singlecell datasets. On the other hand, we evaluate the preservation of local and global structures of the original datasets in the low-dimensional embeddings. In each case, we consider multiple evaluation metrics.

#### Three biology-preservation metrics

For a given low-dimensional embedding and a set of cellspecific biological labels, we consider the following three evaluation metrics. The Silhouette index measures the cohesion and separation of clusters [62]. It is formally defined as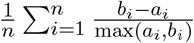where *a*_*i*_ is the average distance to points in the same cluster and *b*_*i*_ is the average distance to points in the nearest cluster. A higher Silhouette Score indicates better alignment between the data points and the cluster labels, with points being closer to their own clusters than to other clusters. The function `silhouette()`in the R package `cluster` is used. The neighbor purity score measures the proportion of a cell’s *k*-nearest neighbors belonging to the same group as that cell. A higher neighbor purity score indicates that the groups are more homogeneous. The function `neighborPurity()` in the R package `bluster`is used, with *k* = 50. The local Simpson score is defined as the reciprocal of the Local Inverse Simpson index (LISI), defined in [39] and implemented as the function `compute _ lisi()` in the R package `lisi`. LISI estimates the effective number of clusters contained in a cell’s neighborhood. As such, a higher local Simpson score indicates better separation of the clusters.

#### Two structure-preservation measures

For a given pair of input high-dimensional dataset and its low-dimensional embedding, we consider the following two evaluation metrics. The correlation score measures the similarity between pairwise distances in the original and reduced embedding spaces, defined as corr (dist(**Y**), dist(**X**)), where dist(**X**) is the pairwise Euclidean distance among cells computed from the high-dimensional dataset and dist(**Y**) is the pairwise Euclidean distance among cells in the low-dimensional embedding. A higher correlation score indicates that the lowdimensional embedding preserves pairwise distances more faithfully. The (neighbor) concordance score quantifies how well the neighborhood structure is preserved between the high-dimensional data (**X**) and the low-dimensional embedding (**Y**). Formally, it is defined as

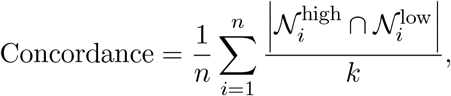

where *n* is the total number of data points in the dataset,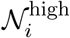 is the set of *k*-nearest neighbors of point *i* in the high-dimensional space,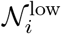 is the set of *k*-nearest neighbors of point *i* in the reduced-dimensional space, and *k* is the number of nearest neighbors considered. A higher concordance score indicates better preservation of neighborhood relationships between the original and reduced-dimensional spaces. We used *k* = 100 in all our analyses.

#### Simulated datasets

We generate simulated datasets containing smooth structures to evaluate the performance of NE algorithms using the metrics introduced above. In the first case, we randomly sampled *n* = 2000 data points from a one-dimensional curve embedded in high-dimensional space. The curve is divided into five equal-length segments, and each sample is assigned a “pseudotime” label (from 1 to 5) based on the segment to which it belongs. In the second case, *n* = 2000 data points were randomly sampled from the unit circle. In both examples, the simulated datasets contain a low-dimensional (smooth) manifold structure.

### 4.2 Theoretical Insights on NE Algorithm Artifacts

To better understand the nature of the empirically observed artifacts in NE algorithms with default parameters, we conduct a rigorous theoretical analysis of t-SNE, the most basic NE algorithm. We formally establish the connection between low GCP values and these artifacts, particularly the fragmentation of the underlying manifold structure, in the final low-dimensional embeddings. We expect our findings about t-SNE to generalize to other NE algorithms, given the similarities in their loss functions and iterative optimization schemes [12, 38, 21].

Recall that the t-SNE algorithm starts by constructing affinities *p*_*ij*_ for each pair of data points by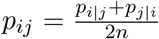, where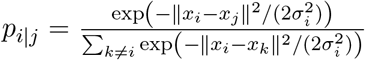, and *σ*_*i*_ ‘s are bandwidth parameters determined through computational procedures such as perplexity quantification detailed in [71]. On the other hand, t-SNE also defines embedding affinities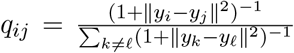, where *y*_*i*_ ‘s are the desirable low-dimensional embeddings. The objective of t-SNE is to minimize the KL-divergence between affinities {*p*_*ij*_} and {*q*_*ij*_}, given by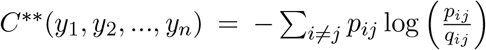. Since the affinities {*p*_*ij*_} are fixed for given data set {*x*_*i*_}_1≤*i*≤*n*_, the above optimization problem is equivalent to solving for {*y*_*i*_} that maximize

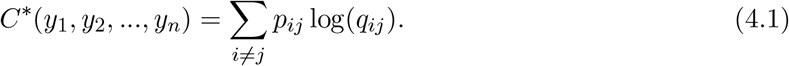

As we argued earlier, an important parameter determining the performance of t-SNE is the graph connectivity characterizing the smoothness of the neighborhood affinity around each point. In the case of t-SNE, the graph connectivity is implicitly determined by bandwith parameter *σ*_*i*_, or “perplexity” in most software implementations. To facilitate our analysis of t-SNE, we consider the following simplification of t-SNE where *σ*_*i*_ = *σ* is constant and therefore the affinities *p*_*ij*_ are approximately given by the *k* nearest neighbors

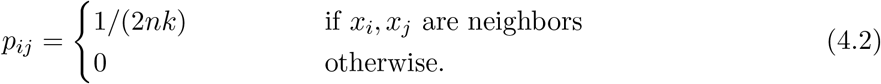

Here the parameter *k* characterizes explicitly the graph connectivity in the affinity matrix *P* = (*p*_*ij*_). To provide deeper insights on the reason behind the empirically observed distance-distortion of NE algorithms for embedding manifold-like structures under lower GCPs, we consider the following <u>prototypical case</u> where t-SNE is applied to embed data points uniformly distributed on a unit circle in ℝ^2^. Despite the simplicity of the example, we believe that our analysis can be generalized to all one-dimensional manifolds. Since this is beyond the scope of the current work, the fully generalized theoretical results will be developed elsewhere in a more systematic manner. Consider the discrete circle of *n* points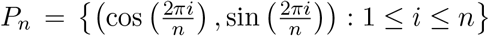in ℝ^2^. Our goal is to prove that, when the graph connectivity parameter is relatively small compared with the sample size *n*, the <u>optimal</u> t-SNE embedding *ϕ* : *P*_*n*_ → ℝ^2^ must distort distances, and the distortion becomes even bigger as *n* increases. The natural mathematical framework that describes metric distortion is that of bilipschitz embeddings. Specifically, a mapping *ϕ* : *X* → *Y* is bilipschitz with distortion parameter *L* ≥ 1 and scaling factor *S >* 0 if

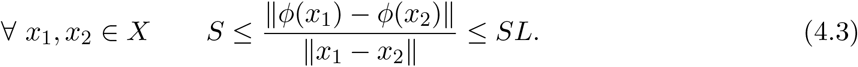

Intuitively, a diverging distortion parameter *L* suggests that the mapping *ϕ* is not Lipschitz, essentially containing points of discontinuity, or “gaps” in its image. The following theorem, whose proof can be found in the Supplement, states that, whenever the nearest neighbor (or graph connectivity) *k* is relatively small compared with *n*, and if the diameter of the low-dimensional embedding is of order *o*(*n*) (which is clearly satisfied empirically in all our examples), the optimal t-SNE embedding given by the solution of (4.1) will have a diverging distortion parameter *L*. This theorem explains the fragmentation patterns empirically observed in the final visualizations.

#### Theorem 4.1

(t-SNE artifact). *Let P*_*n*_ *be the discrete circle and let ϕ*^*^ *be the optimal embedding induced by the t-SNE objective (4.1) with the affinities (4.2) and the graph connectivity parameter k. Suppose S*(*ϕ*^*^) = *n*^*τ*^ *for some* 0 *< τ <* 1 *and k* = *O*(1) *as n* → ∞. *Then, the distortion factor of ϕ*^*^ *is unbounded, that is, L*(*ϕ*^*^) → ∞ *as n* → ∞. *Proof*. See Section B of the Supplement.

### 4.3 ness Approach

In this section, we provide details of the proposed NESS approach. We begin by describing how to generate algorithmic perturbations for any NE algorithm using random initialization, followed by the construction of a KNN matrix that captures embedding stability. We then introduce the definitions of the NESS local stability score, global stability score, and embedding rareness score. Finally, we present a data-driven NESS workflow that automatically identifies the optimal GCP.

NESS is designed to generate stability measures of an NE algorithm for better hyperparameter tuning, interpretation, and downstream inferences. It evaluates the stability of neighborhood graphs derived from the low-dimensional embeddings across multiple NE runs based on different random initializations. First, for a given a dataset and NE algorithm under some hyperparameter configuration, we run *N* times under random initialization to generate *N* low-dimensional embeddings, denoted as *Y* ^(1)^, *Y* ^(2)^, …, *Y* ^(*N*)^. For each embedding, we create a *k*-Nearest Neighbors (KNN) graph, represented as *M* ^(*i*)^, where *M* ^(*i*)^[*j*, 1 : *k*] stores the indices of the *k*-nearest neighbors for point *j*. To quantify how often two points are identified as neighbors across all *N* embeddings, we construct a k-nearest-neighbor matrix **K** = (*K*[*j, l*]). This matrix is initialized as a zero matrix of size *n* × *n*, where *n* is the number of data points. As *i* runs from 1 to *N*, the matrix is updated as *K*[*j, l*] ← *K*[*j, l*] + 1 for all *l* ∈ *M* ^(*i*)^[*j*, 1 : *k*]. As a result, the final *K*[*j, l*] represents the matrix entry of **K** at row *j* and column *l*, which stores the number of times point *j* and point *l* appear as *k*-neighbors across the *N* embeddings. To ensure symmetry in the neighbor relationships, we further symmetrize **K** by **K** ← max(**K, K**^⊤^). See Section 4.6 for the recommended values of *k* and *N* in our implementation.

#### NESS local stability score

The local stability score measures the consistency of neighbors for each data point across all *N* embeddings. For a given point *j*, the local stability score *S*_*j*_ is calculated as:

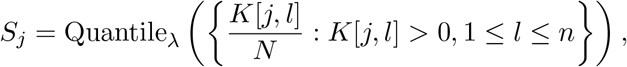

where Quantile_*λ*_ represents the *λ*-th percentile of the normalized neighbor counts. Intuitively, if *S*_*j*_ is large, it means that data point *j* shares many of the same neighbors across the *N* low-dimensional embeddings. In all our analyses we used the recommended value *λ* = 0.75, which is motivated by the expected sparsity of the count matrix **K**. We also found the results to be stable across a wide range *λ* ∈ (0.6, 0.9). The collection of local stability scores is denoted as {*S*_1_, *S*_2_, …, *S*_*n*_}.

#### NESS global stability score

The global stability score provides an overall measure of stability for the dataset by aggregating the local stability scores. It is computed using the mean of the local stability scores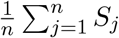 across all cells. A higher Global Stability score indicates that the neighbors of a point are more consistently preserved across different embeddings. To select an appropriate GCP threshold for a given NE algorithm, we recommend choosing the low-dimensional embedding corresponding to the smallest GCP value that falls within the top 5% of all evaluated global stability scores.

#### NESS embedding rareness score

The NESS embedding rareness score quantifies the dissimilarity of each NESS-generated low-dimensional embedding from other *N* − 1 embeddings. We start by counting how many times two cells share the same neighbor in the embeddings. For each pair of embedding *i* and *j*, for each data point *l*, we evaluate the number *w*_*ij,l*_ of shared *k*-nearest neighbors and define the similarity measure *W* (*i, j*) = median({*w*_*ij,l*_*/k* : 1 ≤ *l* ≤ *n*}). Repeat this for all pairs of embeddings, we obtain an *N* × *N* pairwise similarity matrix *W*. For each embedding *i*, we evaluate the mean and variance of the *i*-th column of *W* (excluding the diagonals), denoted as *m*_*i*_ and *v*_*i*_. Since when both quantities are small, there is bigger dissimilarity between the *i*-th embedding and the other embeddings. We define the embedding-specific rareness score *R*_*i*_ = 1*/*(*m*_*i*_*v*_*i*_). By this definition, we hope to highlight (through a bigger embedding rareness score) embeddings which are (i) on average more dissimilar with other embeddings (i.e., with smaller *m*_*i*_), and (ii) more uniformly deviated from other embeddings (i.e., with smaller *v*_*i*_). A higher embedding rareness score indicates that the embeddings differ significantly from the others, likely containing more severe distortions and artifacts.

#### An automated NESS workflow

To improve computational efficiency and practicality of our proposed method, we propose an automated NESS workflow. For a given dataset and NE algorithm, we first consider a sequence of five equally spaced GCP parameters ranging from 10 to 10 log *n*, where *n* is the sample size. Starting from the smallest GCP, we evaluate the NESS global stability (GS) score of the corresponding low-dimensional embedding, and stop if either the GS under the current GCP exceeds 0.9, or the GS under the current GCP is not greater than that under the previous, smaller GCP by at least 5%. The final GCP is chosen as the recommended GCP. Under this GCP, we record and output the associated local stability scores, the NESS embedding rareness score (optional), and the final low-dimensional embedding.

### 4.4 Biological Case Studies to Demonstrate the Effectiveness of NESS

In this section, we first define a local density metric and connect it to the NESS local stability scores obtained in four single-cell datasets, to provide an explanation of the observed local instability of NE algorithms. Then we demonstrate the effectiveness of NESS in three different biological processes in (i) identifying transitional and stable cell states, (ii) quantifying cell-specific transcription activity dynamics, and (iii) differential expression analysis and pathway enrichment analysis.

#### Quantifying local density

For each *p* from 0 to 100, we focus on the cells whose NESS local stability score lies in the upper *p*-th percentile across all cells. Denote the indices of these cells as the set *I*_*p*_. Then for each cell in *I*_*p*_, we compute its mean Euclidean distance to its 30-nearest neighbors in the PCA dimension-reduced feature space. Then we normalize the mean distances by dividing them by the 50% percentile of all the pairwise distances between the 30-nearest neighbors. The normalized distance characterizes the average local density of the cells whose local stability score is in the *p*-th percentile. For each of the single-cell dataset, by evaluating the average local density of the cells corresponding to different percentiles, we found a strong association: cells with lower local stability scores tend to have lower local density, as reflected by greater average distances from their neighboring cells (Figure 3c).

#### Identifying transitional and stable cell states

To identify transitional and stable cell states in the iPSC, Murine Intestinal, and Embryoid Body datasets, we first apply NESS in combination with an NE algorithm–t-SNE for the iPSC and Murine Intestinal datasets, and UMAP for the Embryoid Body dataset. Our choice of NE algorithm is guided by our empirical observations that NESS(t-SNE) generally performs well on moderate-sized datasets, while NESS(UMAP) offers greater computational efficiency for large datasets (e.g., those with more than 20K cells). The lowdimensional embeddings are obtained under the NESS-recommended GCP in each case. For iPSC and Murine Instestinal datasets, under the NESS recommended GCP, we obtain the NESS local stability score and compare the cell-specific local stability scores with the transition entropy scores obtained from MuTrans to generate Figure 4a. In particular, we compare the entropy values of the cells associated with the bottom *p*% (*p* = 2, 10, 20, 100) of the NESS local stability scores (that is, across the *p*% most unstable cells). For the Embryoid Body dataset, we first apply our algorithm to obtain the NESS local stability score for each cell in the dataset, generating the boxplot in Figure S10a. We then focus on the four neuronal subtypes and rerun our algorithm to obtain the low-dimensional embeddings and the local stability scores (Fig 4d). Then we perform standard differential expression analysis between NS-1/2 and NS-3/4 to identify key genes driving the cell state heterogeneity (Fig 4e).

#### Quantifying cell-specific transcription activity dynamics

For both Spermatogenesis and Dentate gyrus neurogenesis datasets, we first applied NESS to determine a suitable hyperparameter configuration of the NE algorithm (UMAP in our analysis), to obtain the corresponding NESS local stability scores {*S*_1_, …, *S*_*n*_}. Then, for each cell *i*, we define the pointwise NESS instability score as {1*/S*_1_, …, 1*/S*_*n*_} (Figure 5). For better visualization and comparison with total RNA velocity, we apply the monotone transformation log(1 + 100 log(1*/S*_*i*_)) to the instability score. Once computed on these datasets, the NESS instability scores are grouped by cell states and then compared with total RNA velocity, which is defined as the *L*_2_-norm of the estimated RNA velocity vector for each cell. In the Spermatogensis dataset, the spermatid pseudotime is estimated by projecting cells onto the differentiation trajectory within the NESS-assisted UMAP embedding.

#### Differential expression analysis and pathway enrichment analysis

For the iPSC dataset, we began by identifying genes whose expression are significantly associated with the NESS(t-SNE) local stability score using Pearson correlation test. As a result, 48 genes were found significant with a BH-adjusted p-value below 0.05 (Table S2). To identify the key biological pathways associated with these significant genes, we use the DAVID Functional Annotation Bioinformatics Microarray Analysis tool (https://davidbioinformatics.nih.gov/). The DAVID platform identified 7 significant KEGG pathways including the Wnt signaling pathway, the Notch signaling pathway, the Hedgehog signaling pathway, the Cytokine-cytokine receptor interaction, the HIF-1 signaling pathway, the TGF-beta signaling pathway, and the MAPK signaling pathway.

### 4.5 Computing Time

The primary computational cost of NESS arises from generating multiple neighbor embeddings of a dataset using different random initializations. To improve efficiency, we first compute the pairwise distance matrix or the k-nearest neighbor (KNN) graph from the input data and reuse this information across all rounds of low-dimensional embedding generation, thereby avoiding redundant computations. To empirically evaluate the computing time of NESS combined with different NE algorithms, we first report its runtime across a range of real and synthetic datasets analyzed in this study, for various NE algorithms and GCP values. We observe a generally linear relationship between GCP and runtime (Fig S5a), highlighting the efficiency gains enabled by our automated GCP selection procedure.

We also compare the runtime of NESS(UMAP) to that of MuTrans, a popular method for modeling cell state transitions, across datasets with varying numbers of cells. While NESS(UMAP) is slightly slower than MuTrans for datasets with fewer than 1, 000 cells, it demonstrates significantly better scalability for larger datasets, particularly those with over 10^4^ cells (Fig S5b). In particular, for datasets of approximately 10^4^ cells, NESS(UMAP) completes in less than 17 minutes, which is the same amount of time MuTrans requires to analyze a dataset of only about 4,000 cells. Thess results demonstrate the computational efficiency of NESS and its scalability to large single-cell datasets as compared with the existing methods.

### 4.6 Implementation Details

#### GCP definition and implementation details of NESS

For all the analyses presented in this study, we utilized the NESS approach, implemented in R, which supports t-SNE, UMAP, and PHATE for embedding high-dimensional data. The algorithm quantifies the stability of lowdimensional representations using k-nearest neighbor (k-NN) graphs. For t-SNE, the parameter `perplexity` is identified as the GCP parameter, and random initialization is used. For UMAP, the algorithm is randomly initialized with `lvrandom` and the nearest neighbor parameter n neighbors is identified as GCP. For PHATE, we apply `phate` with `ndim = 2`, identify `knn` as the GCP, and used a fixed seed for reproducibility. For DensMAP, we identify `n_neighbors` as the GCP. All the other parameters are set as default values.

The function takes as input a set of GCP parameters, pre-processed (such as standard quality control and count normalization pipelines for single-cell data) high-dimensional data, an optional cell type annotation list, and the chosen dimensionality reduction method (`“tsne”, “umap”`, or `“phateR”`). By default and in all our analyses, we set `N = 30` independent runs and used `k = 50` neighbors for k-NN calculations. The stability of embeddings was quantified using a `knn.score` computed over multiple runs, with a stability threshold of `0.75` by default.

#### Data preprocessing

For each dataset analyzed in this study, we first applied the singular value hard-thresholding algorithm (Algorithm 2 of the Supplementary Notes of Ma et al. [51]) to the normalized and scaled count matrices to denoise the data as a preprocessing step. The number of singular values is determined by *r*_*max*_ = max{*r* : 1 ≤ *r* ≤ *n, λ*_*r*_*/λ*_*r*+1_ *>* 1 + *c*} for some small constant *c >* 0 such as 0.01. Theoretical justifications of such a method have been established based on standard concentration argument [82]. For the Dentate gyrus dataset, we excluded the rare cell types/states with less than 50 cells. For the Spermatogenesis dataset, we excluded the cell types/states with less than 5 cells, and renamed “A3-A4-In-B Differentiating spermatogonia” as “spermatogonia”, renamed “Leptotene/Zygotene spermatocytes” as “early spermatocytes”, combined “DIplotene/Secondary spermatocytes” and “Pachytene spermatocytes” into “late spermatocytes”, and combined “Early Round spermatids”, “Mid Round spermatids” and “Late Round spermatids” into “spermatid”. The Murine Intestinal data, Dentate gyrus data, and Spermatogenesis data were pre-processed in previous studies, giving the 2,000 most variable genes.

#### Implementation of baseline methods

Below we describe our implementation of other existing methods.

- **scDEED**: The R function `scDEED`from the scDEED package with `K = 50`, `reduction.method = “tsne”`, and `perplexity`set to the corresponding GCP value. Other parameters were set to default values.
- **pyMuTrans**: The Python function `pm.dynamical_analysis` from the `pyMuTrans` library with `choice distance = “cosine”`, `perplex` set to the corresponding GCP value, `K_cluster = 9.0`, `trials = 10`, and `reduction coord = “tsne”`. Other parameters were set to default values.
- **EMBEDR**: The Python function `calculate_ EES` from the embedr library, using `NearestNeighbors` (from `sklearn.neighbors`) with `n neighbors = 50` to construct a Gaussian kernel-weighted affinity matrix for computing embedding evaluation scores (EES). Other parameters were set to default values.
- **dynamicviz**: The Python function `boot.generate` from the dynamicviz library with `method = “tsne”`, `B = 4`, and `random _seed = 42` for bootstrap sampling. Stability scores were derived using `score.stability_from _variance` with `alpha = 20`. Other parameters were set to default values.
- **scVelo**: The scVelo package was used to preprocess and analyze the RNA velocity dataset. Preprocessing was performed with default parameters. Unspliced and spliced gene expression matrices were smoothed using `scv.pp.moments` with `n_pcs=30` and `n_neighbors=30`. RNA velocity was subsequently inferred using `scv.tl.velocity` and default parameters.

## 4.7 Data Availability

The iPSC data can be downloaded from https://github.com/cliffzhou92/MuTrans-release/blob/main/Data/ipsc.h5ad. The pre-processed Murine Intestinal scEU-seq data can be downloaded from figshare https://doi.org/10.6084/m9.figshare.23737116.v1. The Embryoid Body data can be downloaded from figshare https://doi.org/10.6084/m9.figshare.23737416.v1. The Mouse Hema data can be downloaded from the PhateR GitHub repository https://github.com/KrishnaswamyLab/PHATE/blob/main/data/BMMC_myeloid.csv.gz. The pre-processed Spermatogenesis data can be accessed from the R package scRNAseq under the data name “HermannSpermatogensisData.” The pre-processed Dentate gyrus dataset can be accessed from the Python package scVelo under the data name “scvelo.datasets.dentategyrus.”

## 4.8 Code Availability

The NESS R package and reproducible analysis scripts can be retrieved and downloaded from our online GitHub repository: https://github.com/Cathylixi/NESS.

## Acknowledgments

RM would like to thank Jonas Fischer, Allon Klein, Dmitry Kobak, Zhexuan Liu, Tiffany Tang, Olivier Pourquié, and Yiqiao Zhong for helpful discussions. RM and BY would like to thank Stefan Steinerberger for inspiring discussions on the theoretical analysis of t-SNE.

## A Supplementary Tables and Figures

**Table S1:** Summary of the analyzed single-cell datasets.

| Data Name | Cell Number | Feature Number | Biological Process | Technology | Citation |
| --- | --- | --- | --- | --- | --- |
| Mouse Hema | 2730 | 3451 | Blood Cell Differentiation | scRNA-Seq | [59] |
| iPSC | 1081 | 96 | iPSC Differentiation | scRT-qPCR | [5] |
| Murine Intestinal | 3452 | 2000 | Murine intestinal organoids development | scEU-Seq | [6] |
| Embryoid Body | 31029 | 19122 | Embryonic stem cell differentiation | scRNA-Seq | [55] |
| Dentate Gyrus | 2930 | 2000 | Dentate gyrus neurogenesis | scRNA-Seq | [31] |
| Spermatogenesis | 2325 | 2000 | Spermatogenesis | scRNA-Seq | [29] |

**Figure S1:**
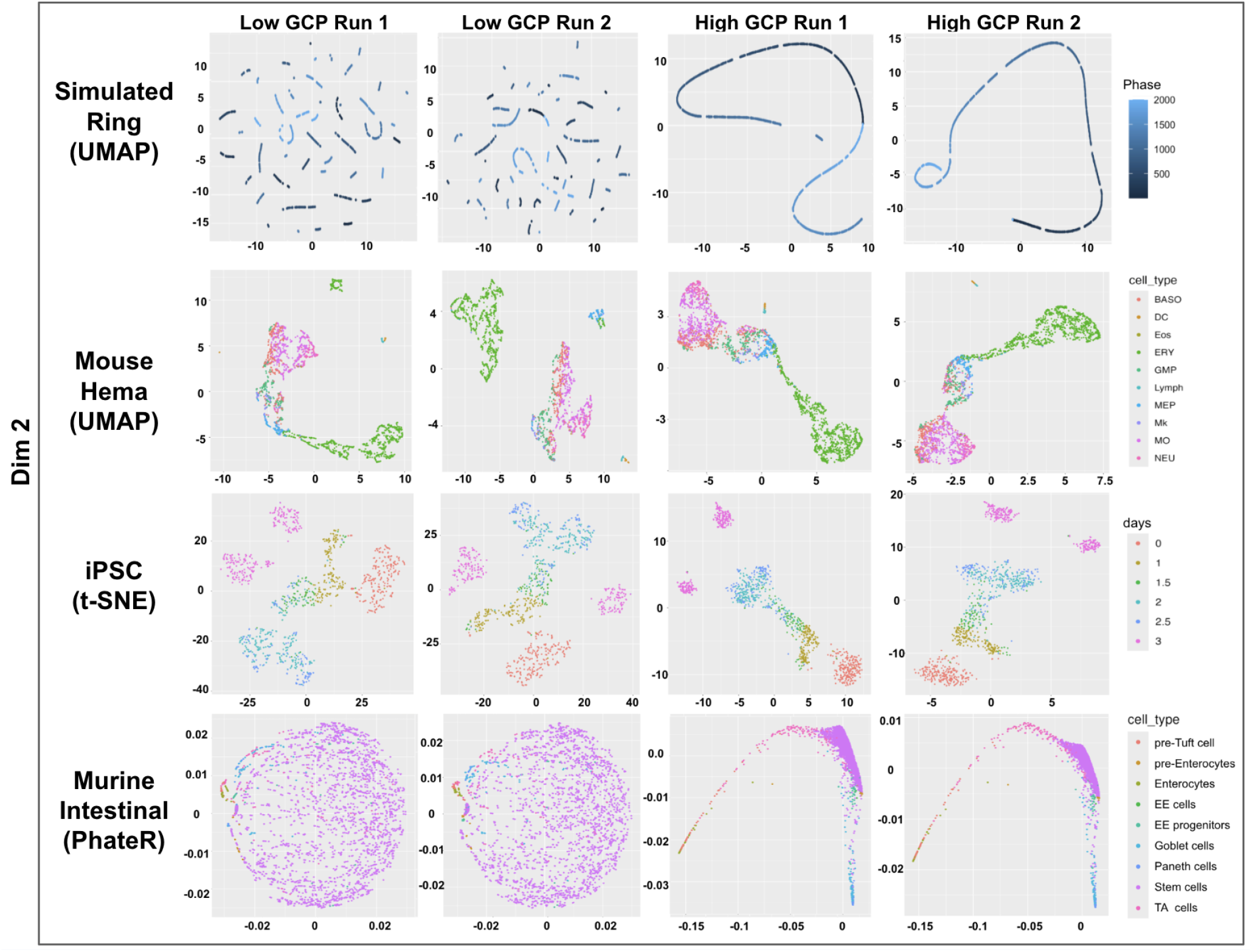
Additional examples of NE embeddings for simulated (first row) and benchmark singlecell datasets (last three rows) generated by popular NE algorithms, with cells colored by their labels. All embeddings are generated using random initialization. The left two columns correspond to two instances of RIs under relative low GCPs, whereas the right two columns show embeddings under relatively high GCP values. The results suggest low GCP can introduce artifacts.

**Table S1:**
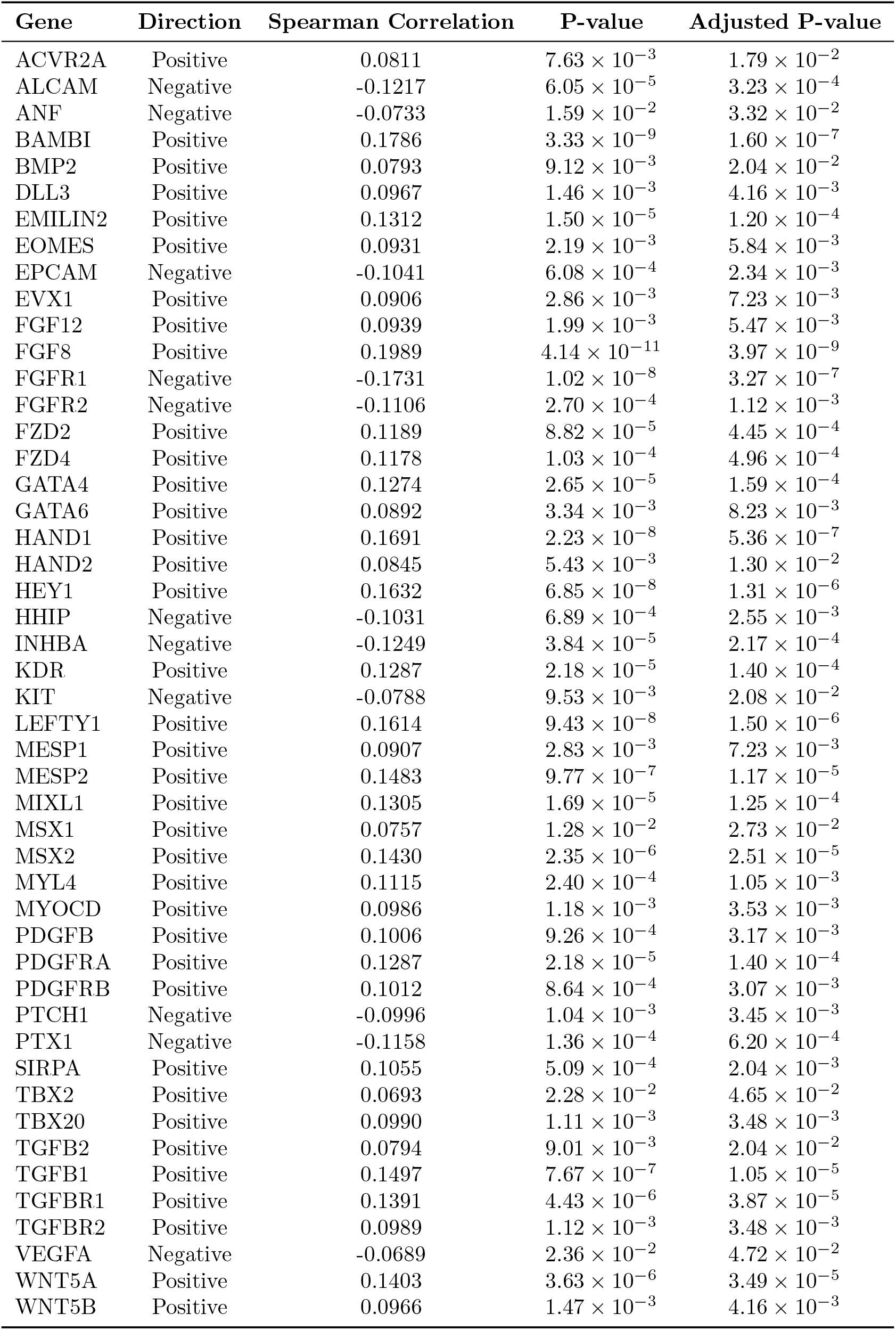
List of key genes in iPSCs with significant correlations (BH-adjusted *p*-value *<* 0.05) between gene expression levels and NESS local stability from t-SNE embedding (GCP = 150, recommended by NESS). The table includes the correlation direction, Spearman correlation coefficients, *p*-values, and BH-adjusted *p*-values. 35

**Figure S2:**
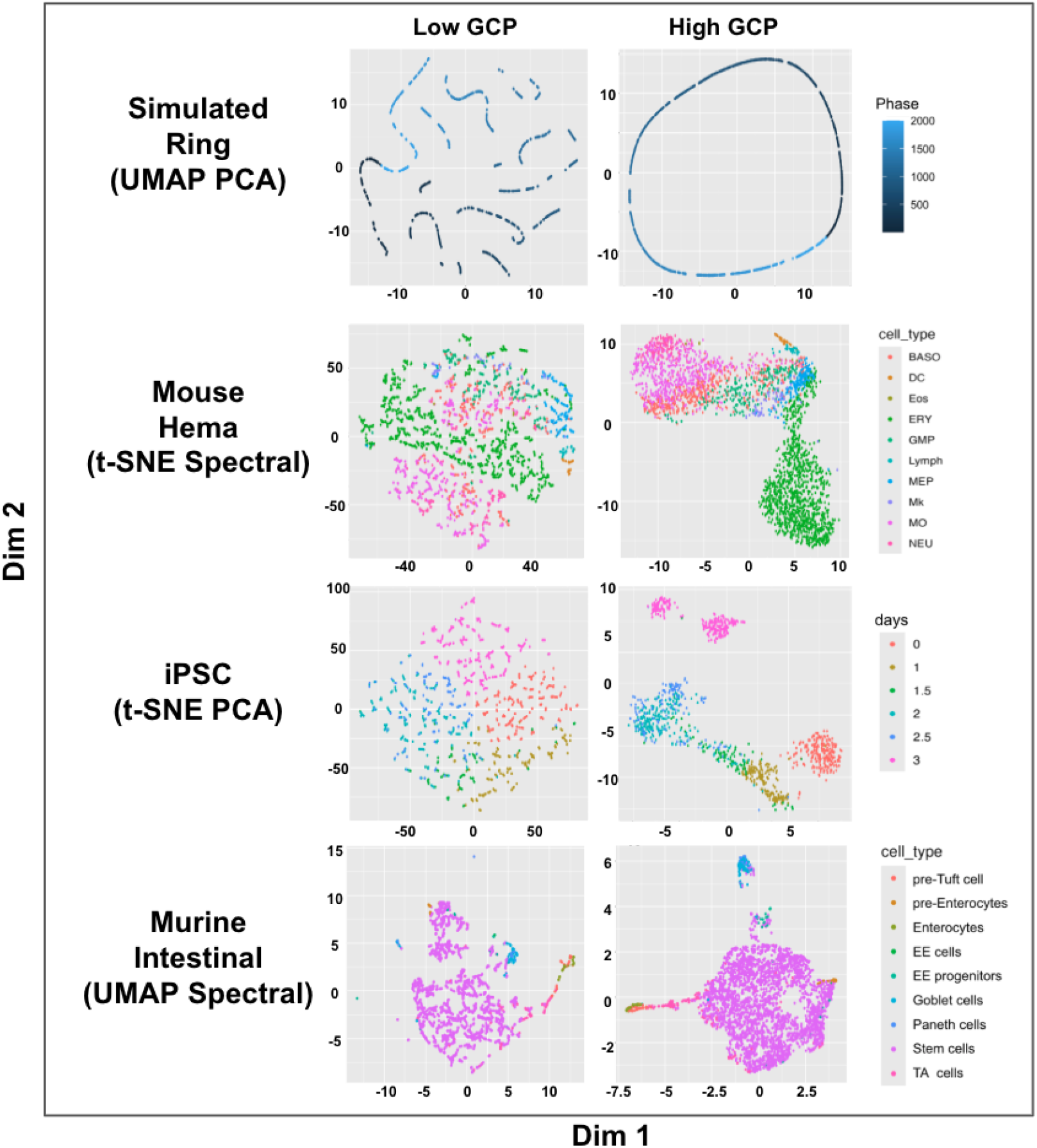
Additional examples of NE embeddings for simulated (first row) and benchmark singlecell datasets (last three rows) generated by popular NE algorithms, with cells colored by their labels. Two embeddings were generated using PCA initialization (first and third rows), and two using spectral (i.e., Laplacian eigenmap) initialization (second and fourth rows). The left column displays representative embeddings under low GCP value, while the right column shows results under high GCP value. The results suggest similar artifacts under low GCP present under other initializations.

**Figure S3:**
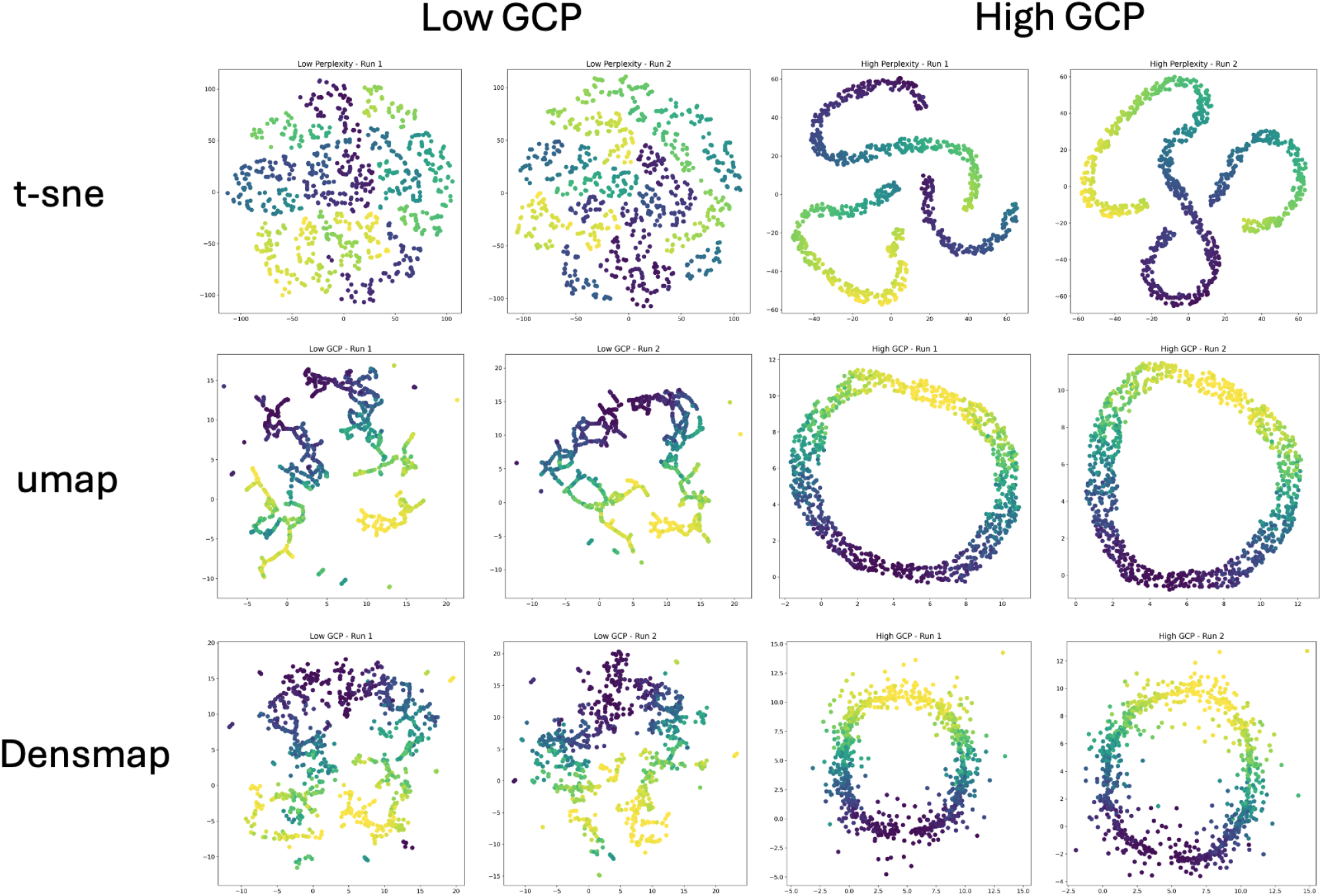
Additional examples of NE embeddings for simulated data (a ring) generated by popular NE algorithms: t-SNE, UMAP, and DensMAP. Data points are colored by their simulated temporal order. Each method was run twice using different random initializations. The left two columns show embeddings generated with low GCP values, while the right two columns show those with high GCP values. The results suggest lower GCP can introduce greater artifacts.

**Figure S4:**
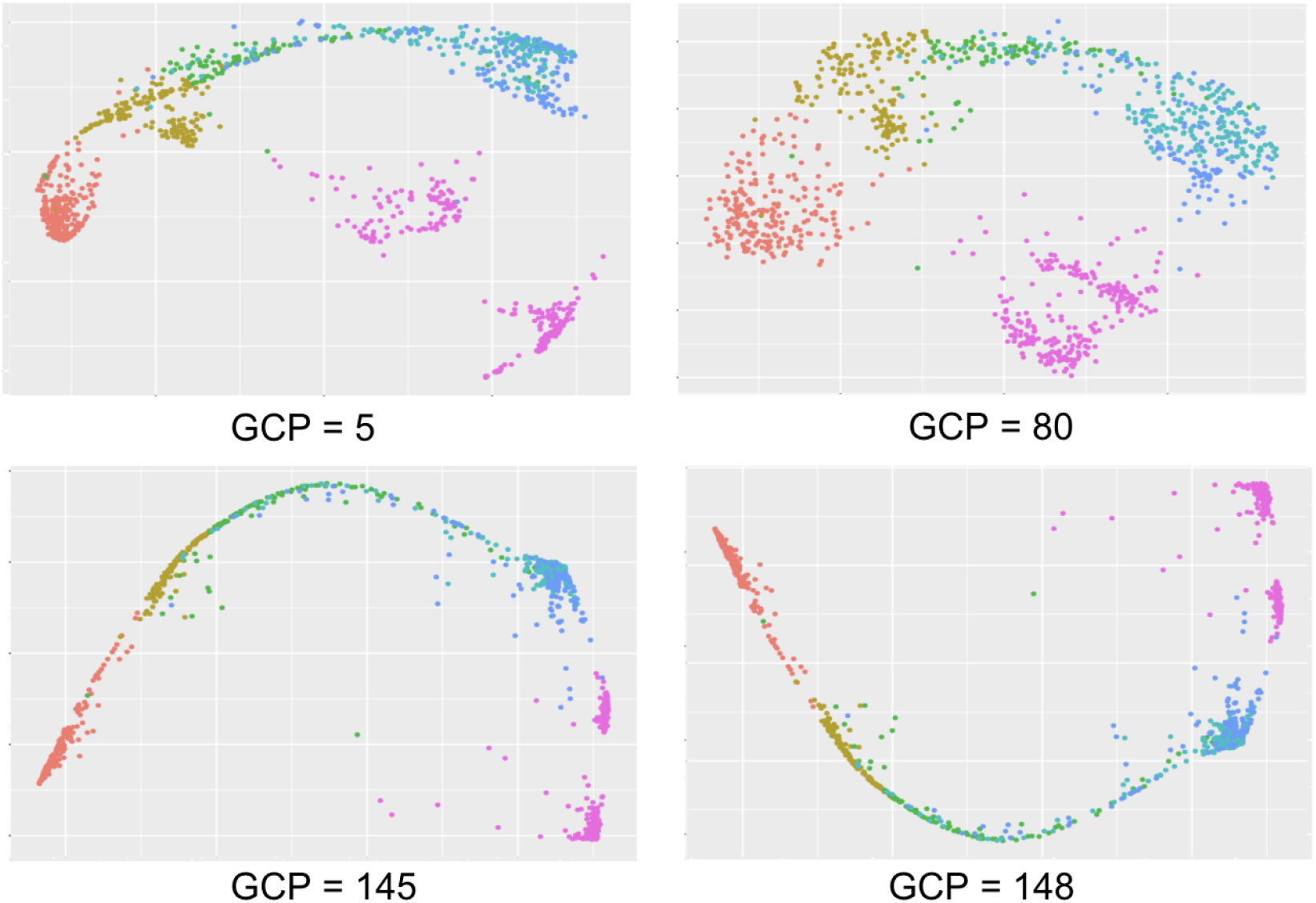
Additional examples of PhateR embeddings of iPSC dataset under different GCP values (knn). For sufficiently GCPs, PhateR likely introduces an over-smoothing effect, causing data points to concentrate along certain learned trajectories while preserving the underlying structure.

**Figure S5:**
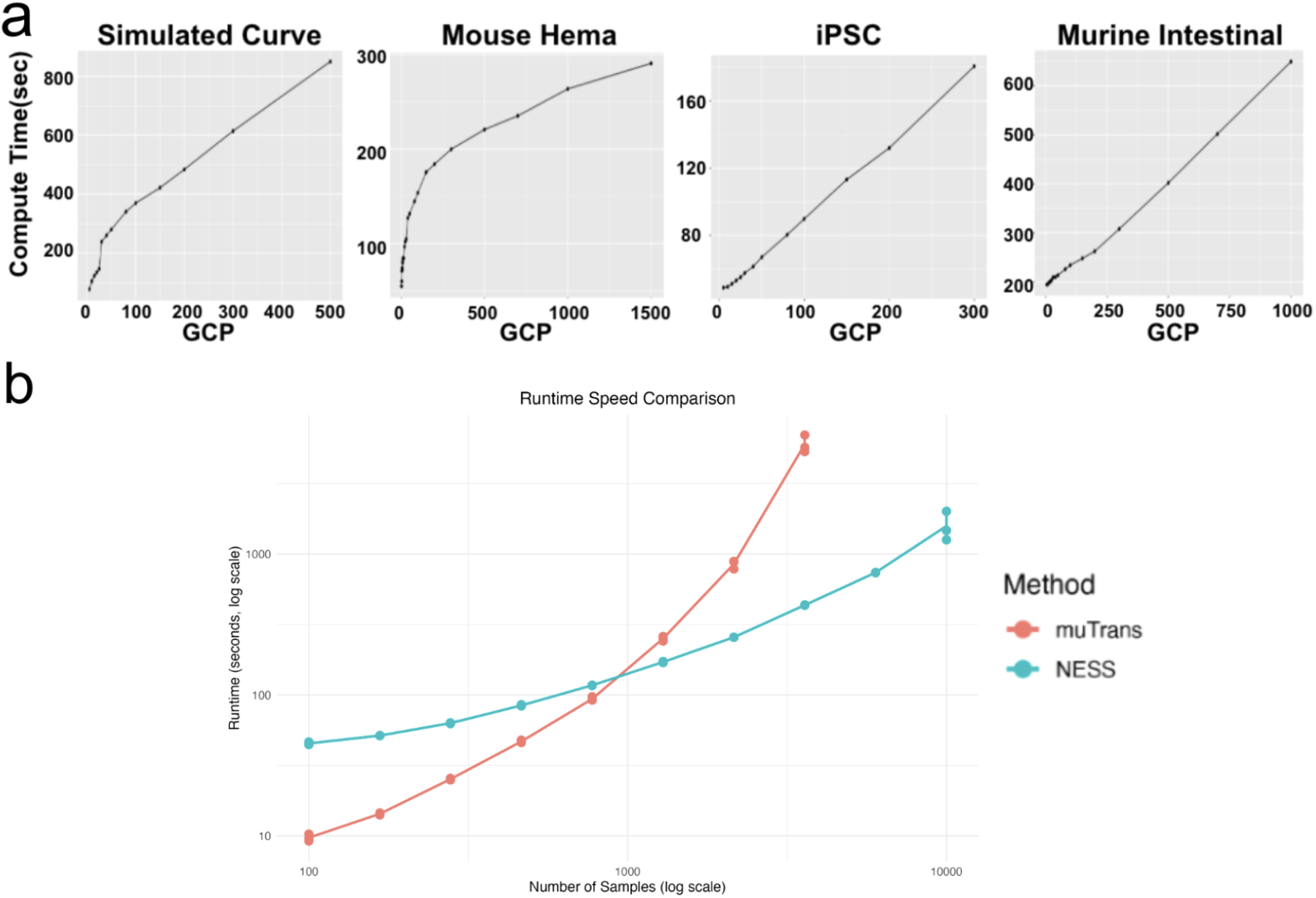
(a) Computation time (in seconds) of the NESS method as a function of the GCP parameter for real and synthetic datasets: Simulated Curve (UMAP), Mouse Hematopoiesis (UMAP), iPSC (t-SNE), and Murine Intestinal (PhateR). A generally linear relationship between GCP and computation time is observed among these examples. All embeddings are initialized with random initialization. In general, a bigger GCP requires more computational time. (b) Runtime comparison between NESS(UMAP) with NESS recommended GCP (=30) and the MuTrans method across various sample sizes. Subsamples ranging from 100 to 10,000 data points were randomly sampled from the Mouse Hematopoiesis dataset to evaluate scalability. This comparison suggests better scalability of NESS to large datasets as compared with MuTrans.

**Figure S6:**
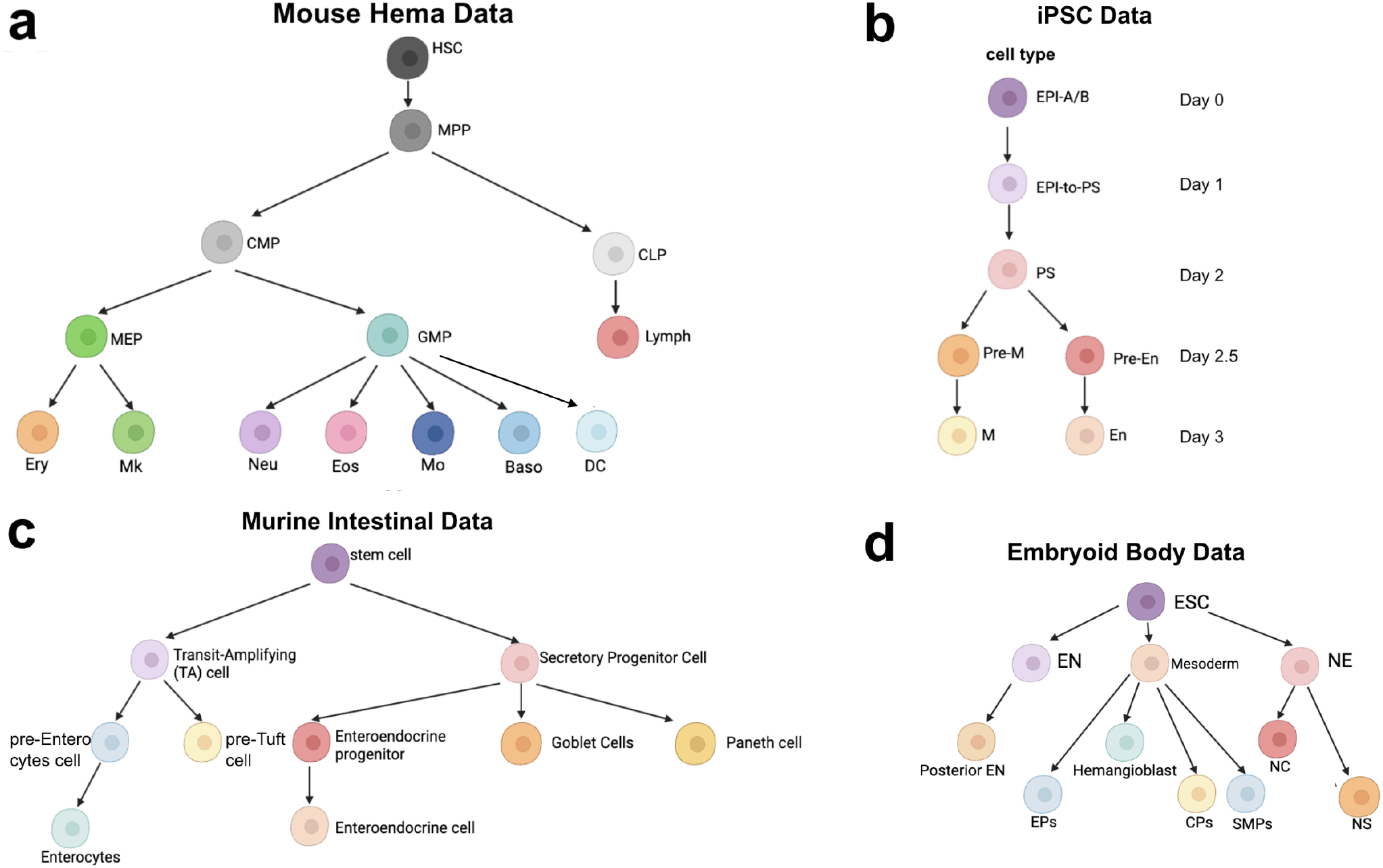
Cell differentiation hierarchies of the benchmark single-cell datasets. (a) Mouse Hematopoiesis dataset [59]. (b) iPSC dataset [5]. EPI: epiblast; PS: primitive streak; M: mesoderm; En: endoderm. (c) Murine Intestinal dataset [6]. (d) Embryoid Body dataset [55].

**Figure S7:**
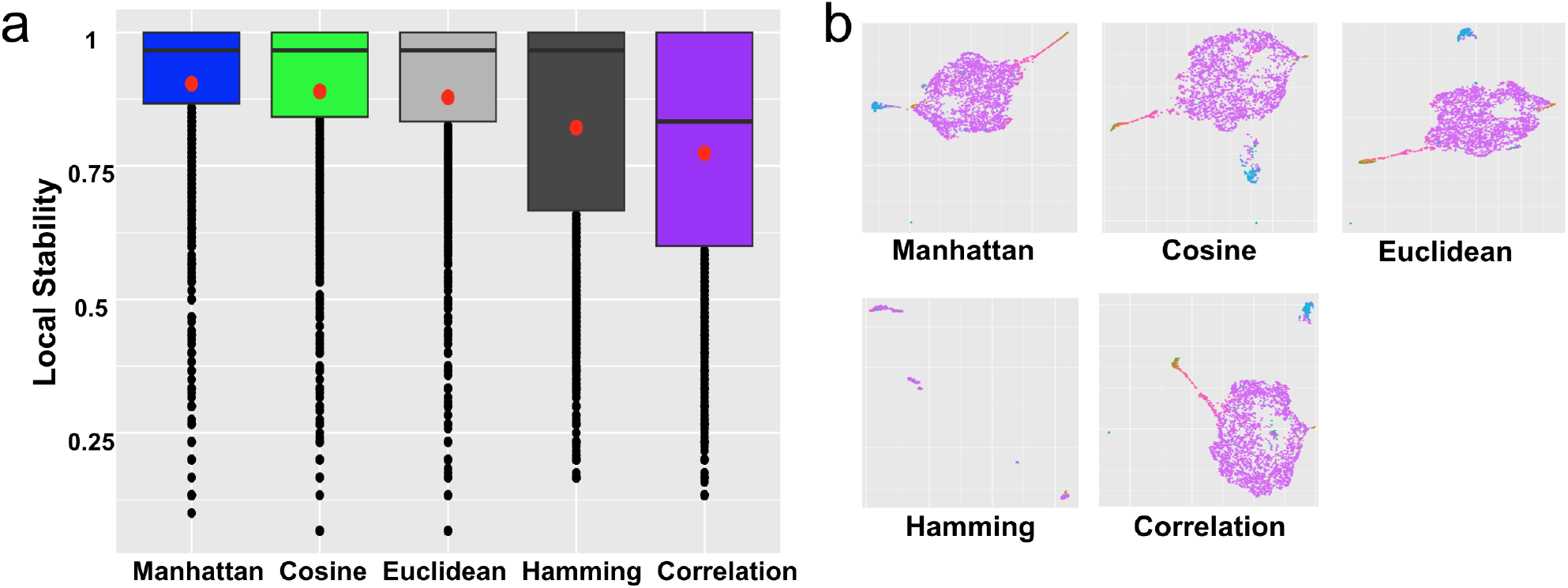
(a) Boxplots of NESS local stability scores for UMAP (GCP=30, recommended by NESS) applied to the Murine Intestinal dataset under different distance metrics. The red dots indicate the corresponding global stability scores. (b) UMAP embeddings (GCP = 30) generated using different distance metrics, colored by cell type. UMAP under Manhattan distance generated the most stable embedding compared with other distance metrics. This result suggests that the NESS global and local stability scores can be used to guide the selection of an appropriate distance metric.

**Figure S8:**
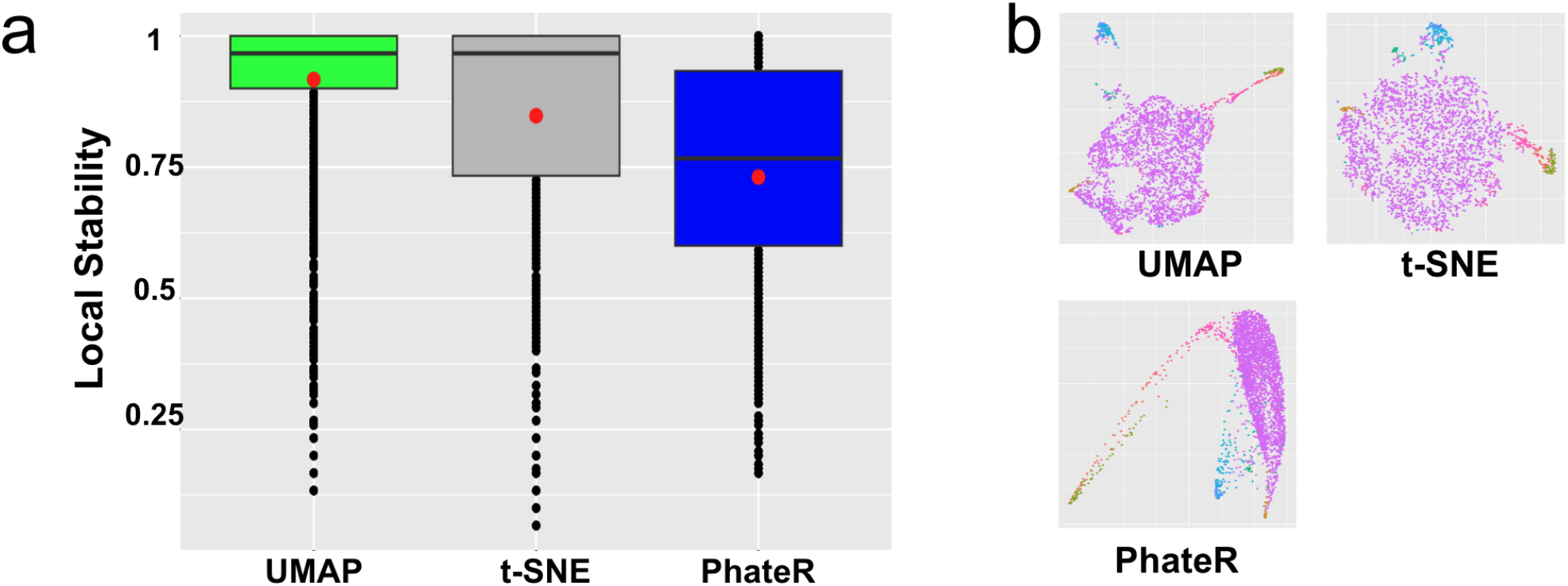
(a) Boxplots of NESS local stability scores for different NE algorithms with GCPs recommended by NESS and other hyperparameters under default setting, applied to the Murine Intestinal dataset. The red dots indicate the corresponding global stability scores. (b) Low-dimensional embeddings generated by different NE algorithms with NESS recommended GCPs, with the cells colored by cell types. UMAP generated the most stable embedding among the three method. This result suggests that the NESS global and local stability scores can be used to guide the selection of an appropriate NE algorithm.

**Figure S9:**
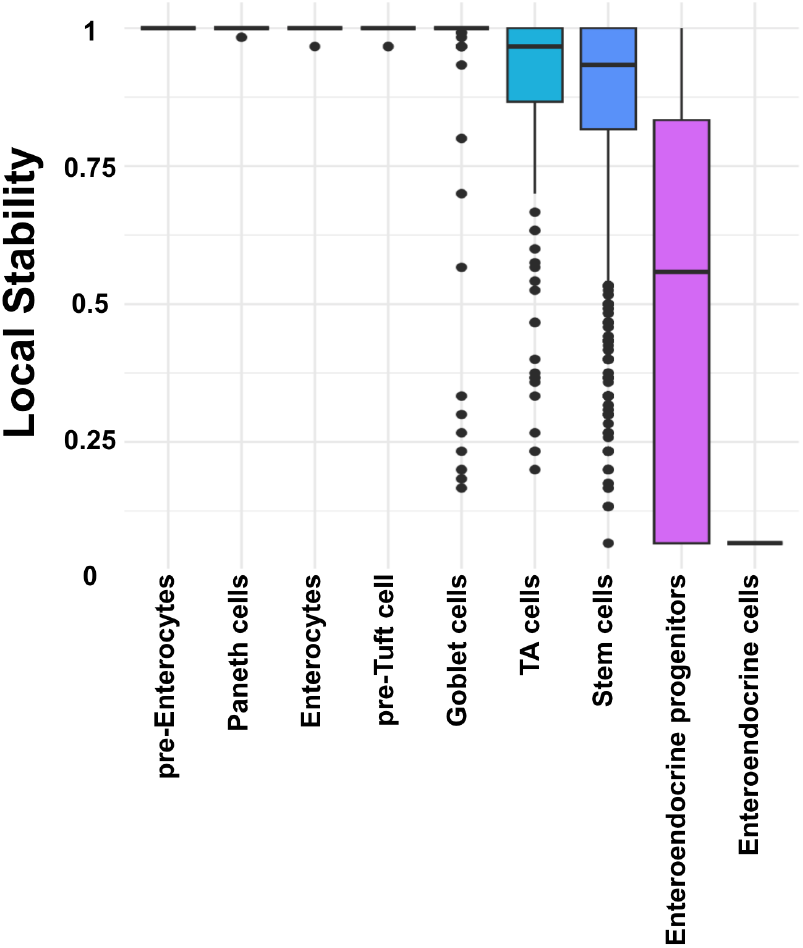
Boxplots of NESS(UMAP) local stability scores for the Murine Intestinal dataset, grouped by different cell types. The stability scores were obtained under the NESS recommended GCP. This result suggests that the NESS local stability score can vary significantly between cell types.

**Figure S10:**
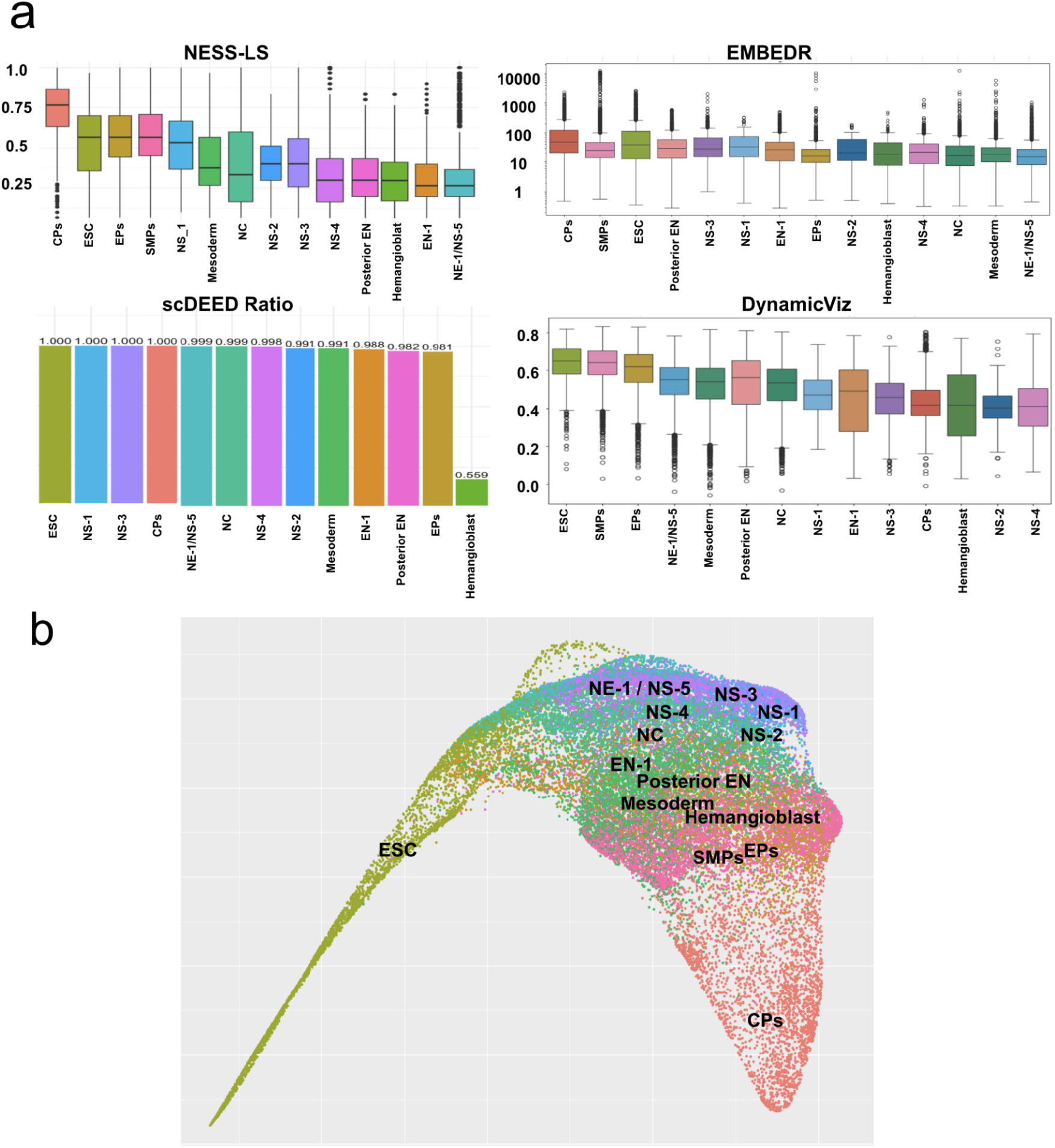
(a) Comparison of cell-type-specific stability scores in the Embryoid Body dataset, ordered by median values. The top-left panel presents NESS local stability (LS) scores, representing local stability estimates for different cell types. The top-right panel shows EMBEDR stability scores, which quantify embedding reliability. The bottom-left panel displays scDEED stability ratios (computed as Trustworthy / (Trustworthy + Dubious)), where lower values indicate a higher fraction of dubious cells. The bottom-right panel illustrates DynamicViz stability scores, capturing variability in the embedding structure. Abbreviated cell type labels include: embryonic stem cells (ESCs), the primitive streak (PS), mesoderm (ME), endoderm (EN), neuroectoderm (NE), neural crest (NC), neural progenitors (NPs), epicardial precursors (EPs), smooth muscle precursors (SMPs), cardiac precursors (CPs), and neuronal subtypes (NS). The results indicates the advantages of NESS in identifying transitional and stable cell states, compared with alternative methods. (b) NESS(PhateR) visualization of the Embryoid Body dataset, with NESS recommended GCP=100. Specific cell types/states are differently colored and labeled on the low-dimensional embedding for better identification.

**Figure S11:**
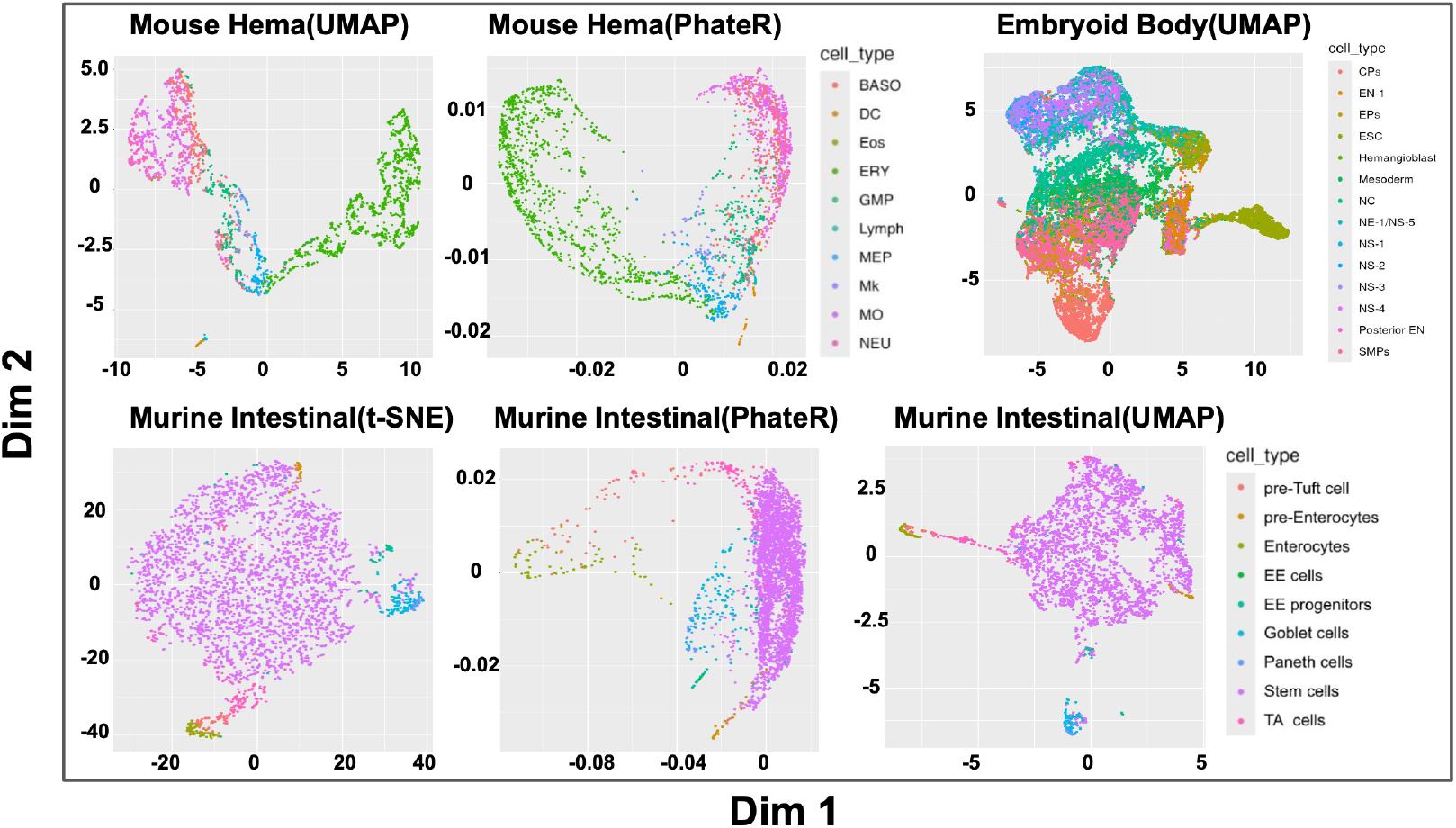
Additional examples of low-dimensional embeddings of single-cell datasets, produced by NE algorithms under default GCPs. For UMAP, the default GCP is n neighbor=15; for t-SNE, the default GCP is perplexity=30; and for PhateR, the default GCP is knn=5. In some cases, the default application of NE algorithm introduces artifacts, such as UMAP’s fragmentation of ERY population in the Mouse Hema dataset. In other cases, the default application of NE algorithms appear to successfully reveal the underlying lineage structure, as observed in the t-SNE and UMAP embeddings of Murine Intestinal dataset.

## B Proof of Theorem 4.1

We start by sketching the proof idea. First, we consider a special mapping *ϕ*_0_ : *P*_*n*_ → ℝ^2^. We show that for any *S*_0_ *>* 0, there exists a specific mapping *Y*_0_ = *ϕ*_0_(*X*) satisfying (4.3) for some *S*(*ϕ*_0_) = *S*_0_ and *L*(*ϕ*_0_) = *L*_0_. In particular, we obtain a lower bound estimate

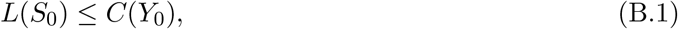

for some function *L*(·) that only depends on *X*. Next, let *Y* ^*^ = *ϕ*^*^(*X*) be the maximizer of *C*(*Y*). We show that, as long as *ϕ*^*^ satisfies (4.3) for some *S*(*ϕ*^*^) ≡ *S*^*^ *>* 0 and *L*(*ϕ*^*^) ≡ *L*^*^ ≥ 1, it holds that

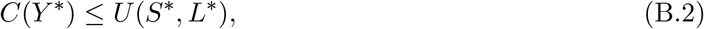

for some function *U* (·, ·) that only depends on *X*. Then it follows that, by choosing *S*_0_ = *S*^*^, we can construct *ϕ*_0_ so that, by (B.2) and (B.1), along with the definition of *Y*\*, it holds that

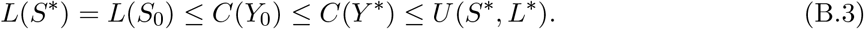

Analyzing the inequality *L*(*S*^*^) ≤ *U* (*S*^*^, *L*^*^) implies a lower bound for *L*^*^. As a result, we obtain that *L*^*^ → ∞ as long as *S*^*^ = *S*(*ϕ*^*^) = *n*^*τ*^ for 0 *< τ <* 1.

### B.1 Technical preparations and additional notation

Note that under the affinities (4.2), the equivalent t-SNE objective (4.1) can be further reduced to maximizing the following objective

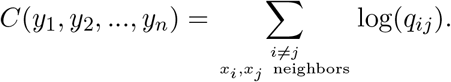

Hereafter, we also denote *Y* = {*y*_*i*_}_1≤*i*≤*n*_ and *X* = {*x*_*i*_}_1≤*i*≤*n*_.

Next we introduce a few useful lemmas. The first lemma collect some elementary mathematical facts which can be found in standard textbooks on analytical geometry and algebra.

#### Lemma B.1.

*For any two points a and b on the unit circle, if the arc distance between a and b is θ, then the length of the chord between them is* 2 sin(*θ/*2).

The next few lemmas concerns upper and/or lower bound estimates of the t-SNE related quantities.

#### Lemma B.2.

*For S* ≳ *n and* {*x*_*i*_}_1≤*i*≤*n*_ = *P*_*n*_, *it holds that*

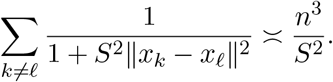

*Proof*. On the one hand, by Lemma B.1, for sufficiently large *n*, we have

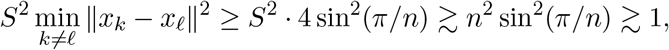

where in the last inequality we use the fact that

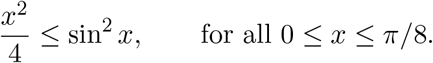

As a result, it follows that

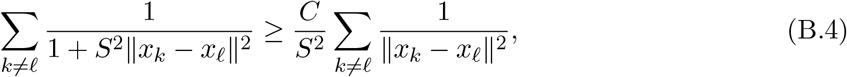

for some absolute constant *C >* 0. On the other hand, we also have

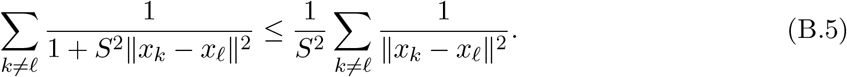

It then holds that

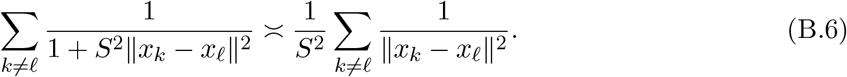

Without loss of generality, we assume *n* is an odd integer. Now we calculate that

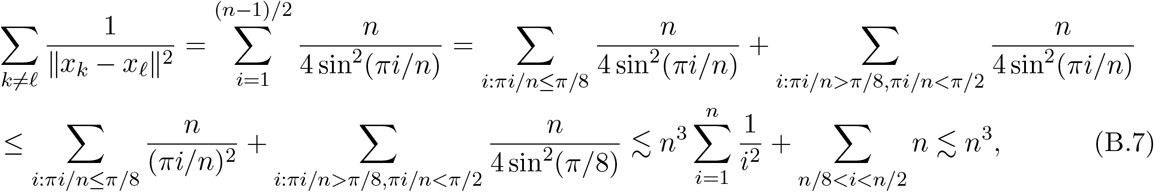

where the last inequality follows from the elementary inequality

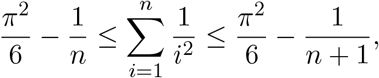

and that

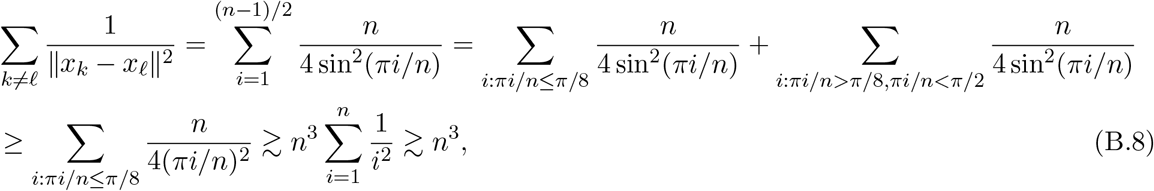

where the third last inequality follows from

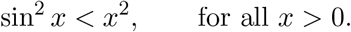

It then follows that

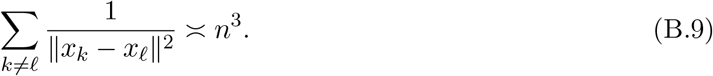

This completes the proof.

#### Lemma B.3.

*Suppose x*_*i*_ *and x*_*j*_ *are k*−*nearest neighbors on P*_*n*_. *Then for any mapping ϕ satisfying (4.3) with the scaling factor S >* 0 *and the distortion factor L* ≥ 1, *for sufficiently large n, it holds that*

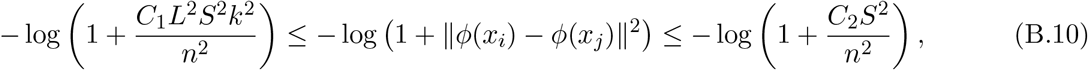

*where C*_1_, *C*_2_ *>* 0 *are some universal constant. In addition, it also holds that*

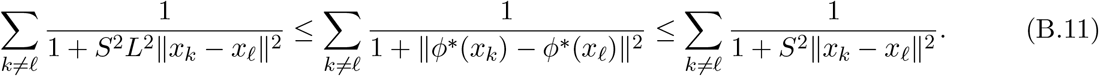

*Proof*. Since *x*_*i*_ and *x*_*j*_ are *k*−nearest neighbors, we have

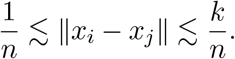

Therefore, by (4.3), we have

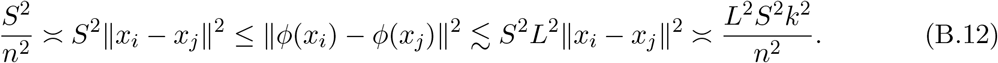

This completes the proof.

**A special mapping** *ϕ*_0_. An important ingredient of our proof is a carefully constructed mapping *Y*_0_ = *ϕ*_0_(*X*_0_), which plays an role in both Part I and Part II of our proof. Below we formally define such a mapping and collect some useful notation and estimates.

Consider the following map *ϕ*_0_ : *P*_*n*_ → ℝ^2^. Without loss of generality, we assume *n* is a multiple of an integer *M*, where

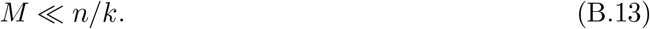

Define a partition 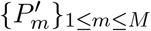of *P*_*n*_ where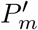 includes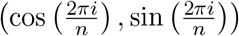for *n/M* consecutive integers *i*. Suppose on each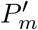, 1 ≤ *m* ≤ *M*, the map *ϕ*_0_ : 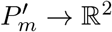 is isometric up to a universal scaling factor *S*_0_, that is,

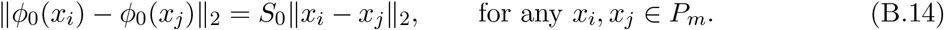

Moreover, we assume that for any *P*_*k*_ ≠*P*_*j*_, there exists an universal constant *c >* 0 such that

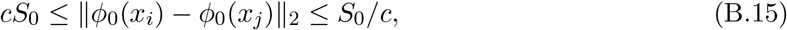

for any *x*_*i*_ ∈ *P*_*k*_ and *x*_*j*_ ∈ *P*_*j*_. The existence of such a map can be established by the following construction: (i) cut the unit circle evenly into *M* parts with equal length; (ii) find the middle point of each curve, connect them into a *M* -lateral shape *Q*; (iii) magnify *Q* by a factor of *M* in all directions; (iv) move the curves isometrically to the corresponding edges of *Q*, without any rotation; (v) scale every point by a factor of *S*_0_.

Before we state our lower bound result, we introduce some useful notation. We define an equivalence relationship such that we write *x*_*i*_ ∼ *x*_*j*_ if and only if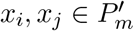for some 1 ≤ *m* ≤ *M*. Now for each 1 ≤ *m* ≤ *M*, we consider a subset *T*_*m*_ of {(*k, 𝓁*) : *x*_*k*_ ≁ *x*_*𝓁*_}, concerning all the points on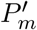 and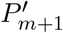 near the edge of the partition point. Note that here we identify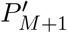 with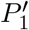. More specifically, if there exists a partition point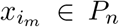 in the sense that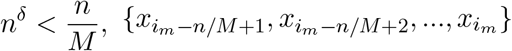and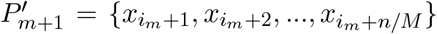, then the subset *T*_*m*_ is defined as

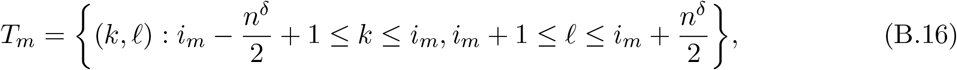

where *δ* ∈ (0, 1) so that *n*^*δ*^ is an even integer, and

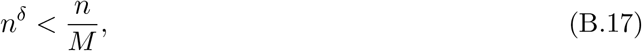

or *M < n*^1−*δ*^. In this way, we have so that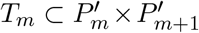and *T*_*𝓁*_ ∩*T*_*k*_ = ∅ for any 1 ≤ *k*≠ *𝓁* ≤ *M*. Apparently, we have

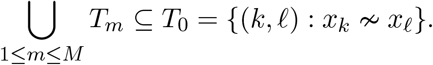

Throughout, we assume

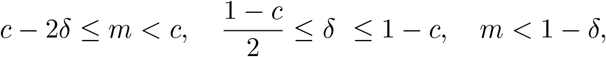

where we calibrate *M* ≍ *n*^*m*^ for some *m* ≥ 0. The following proposition provides a lower bound estimate of *C*(*Y*_0_).

**Proposition B.4**. *Suppose Y*_0_ = *ϕ*_0_(*X*) *is the mapping satisfying (B.14) and (B.15) where S*_0_ = *n*^*c*^ *for some constant* 0 *< c <* 1. *We define T*_*m*_ *as in (B.16) for* 1 ≤ *m* ≤ *M such that (B.13) and (B.17) hold. Then we have*

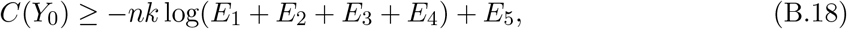

*where*

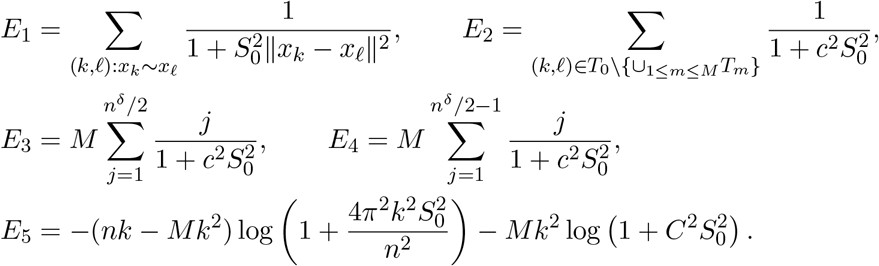

*Proof*. Note that in this case we have

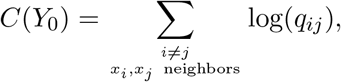

where

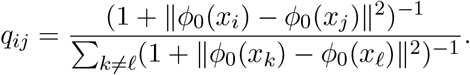

We start from the equation

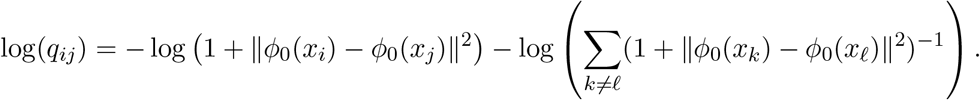

Suppose *x*_*i*_ and *x*_*j*_ are *k*−nearest neighbors. Then we have

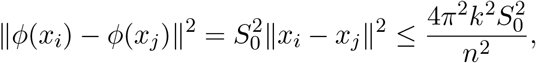

if *x*_*i*_ ∼ *x*_*j*_; and, for *C* = 1*/c*, we have

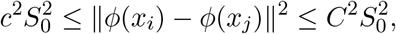

if *x*_*i*_ ≁ *x*_*j*_. As a result, we have

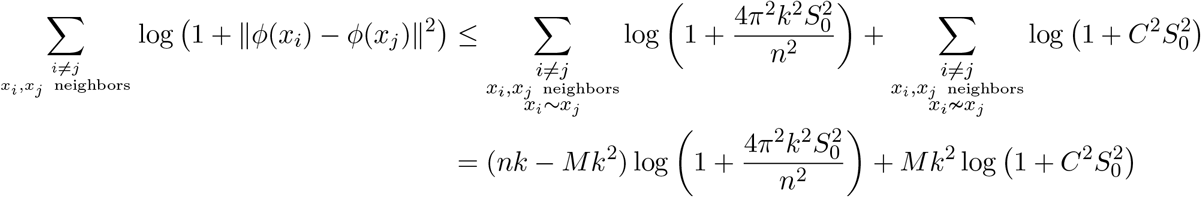

for some constant *C >* 0. On the other hand, we have

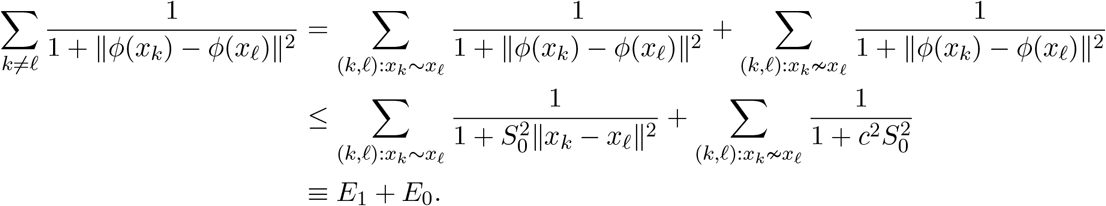

In particular, we have

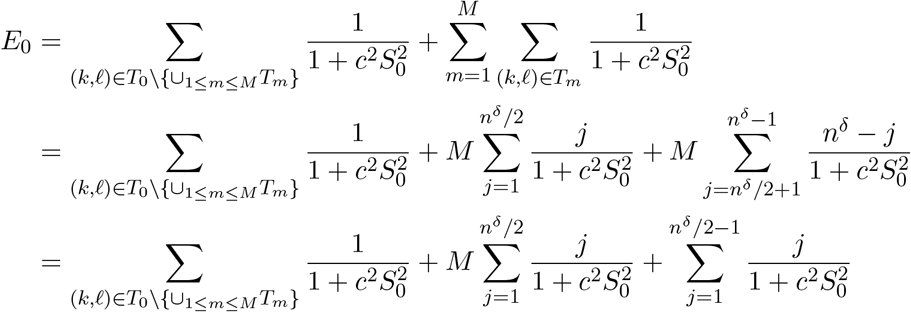

Thus

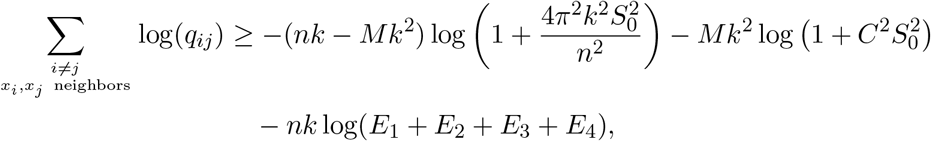

where

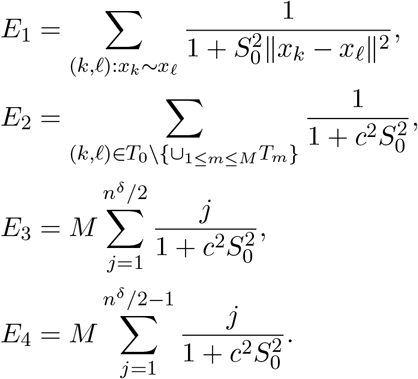

As a result, we conclude that

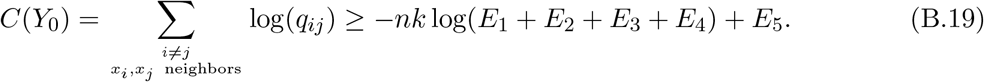

This completes the proof.

#### Proposition B.5.

*Under the assumptions of Proposition B.4, if in addition T*_*m*_ *is chosen such that M* ≲ *n*^*c*^, *it holds that*

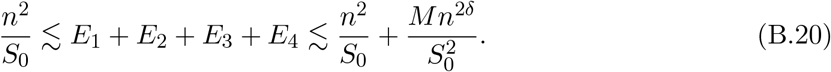

*Proof*. Recall that

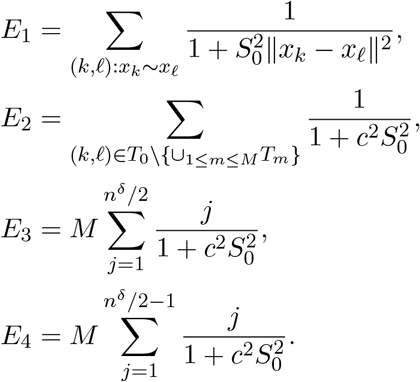

As a result, since *S*_0_ → ∞ as *n* → ∞, we have

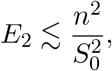

and by the assumption of the proposition, we have

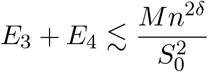

Now for *E*_1_, we consider the following estimates. On the one hand, we consider

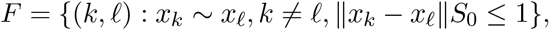

and it follows that

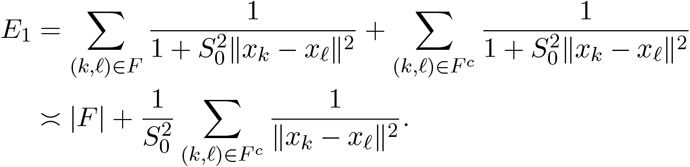

By Lemma B.6, since min_k≠*𝓁*_ ∥*x*_*k*_ − *x*_*𝓁*_∥ ≍ 1*/n* and since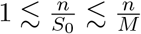, we have

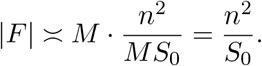

On the other hand, we have

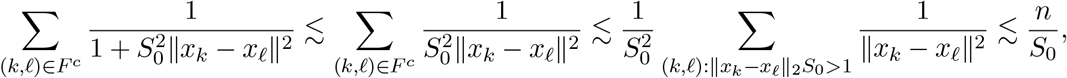

where the last inequality follows from

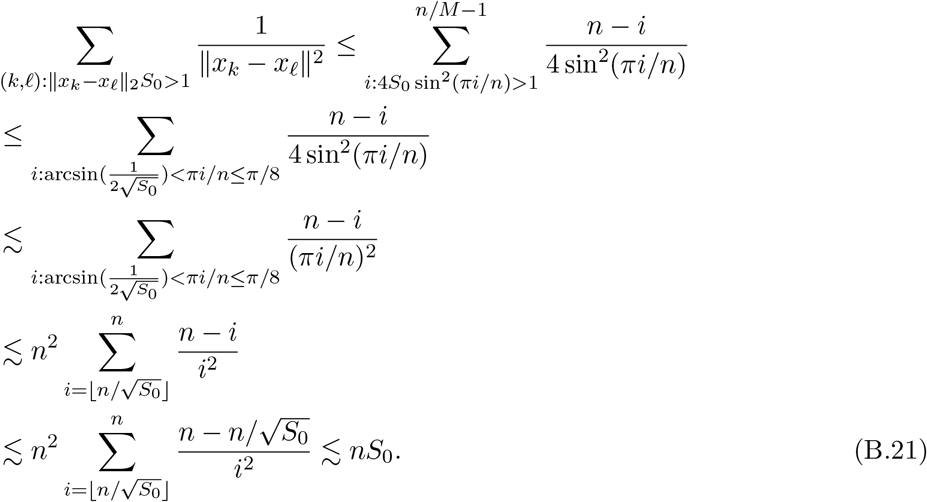

It then holds that

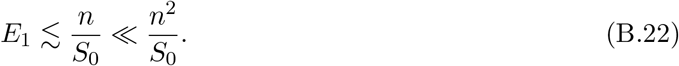

This along with the previous estimates leads to 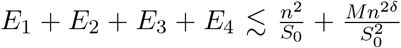. hand, we have

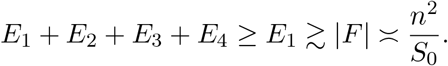

This completes the proof of the proposition.

#### Lemma B.6.

*Let* {*a*_*i*_}_1≤*i*≤*n*_ *be equi-spaced point on* ℝ *so that a*_*i*+1_ − *a*_*i*_ = *h. Let F* = {(*k, 𝓁*) : *a*_*k*_ ∼ *a*_*𝓁*_, |*a*_*k*_ − *a*_*𝓁*_|*S* ≤ 1}. *If* 1 ≪ (*hS*)^−1^ ≪ *n*, then we have 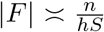. *If instead* (*hS*)^−1^ ≳ *n, then we have* |*F* | ≍ *n*^2^.

*Proof*. Consider a *n* × *n* matrix, whose entries are pairwise distance between {*a*_*i*_}. Let *q* be the biggest integer such that |*a*_1_ − *a*_1+*q*_|*S* ≤ 1. Then we have

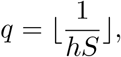

and, if 1 ≤ *q* ≪ *n*,

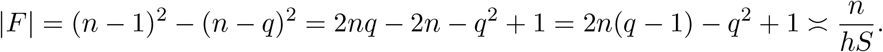

When *q* ≳ *n*, we have |*F* | ≍ *n*^2^. This completes the proof.

The following proposition provides an upper bound estimate of *C*(*Y* ^*^).

**Proposition B.7**. *Suppose ϕ* : *P*_*n*_ → ℝ^2^ *is a bilipschitz embedding at some scale, meaning that there exists a scaling factor S >* 0 *and a distortion factor L* ≥ 1 *such that*

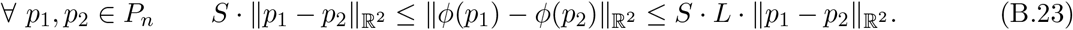

*Moreover, let*

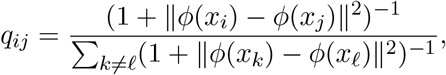

*let* 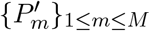*be a partition of P*_*n*_ *satisfying (B.14) and (B.15), and let* {*T*_*m*_}_1≤*m*≤*M*_ *be defined as above. Then*

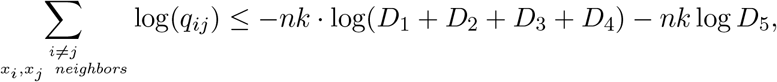

*where*

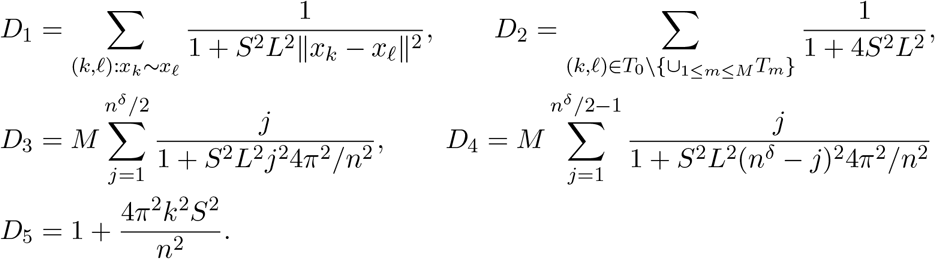

*Proof*. We argue with a pointwise bound. Note that

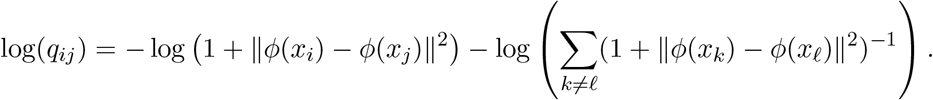

Suppose *x*_*i*_ and *x*_*j*_ are *k*−nearest neighbors. Then we have that

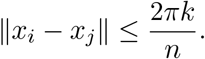

Therefore, by (B.23), we have

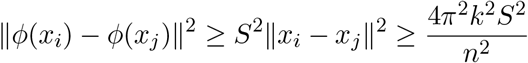

and thus

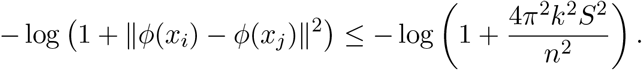

Now we define an equivalence relationship such that we write *x*_*i*_ ∼ *x*_*j*_ if and only if 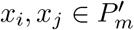 for some 1 ≤ *m* ≤ *M*. We also have

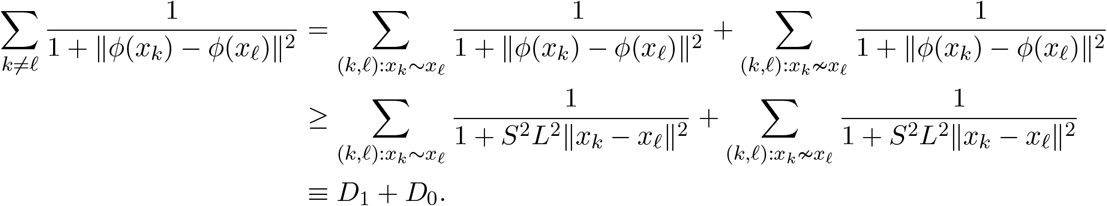

In the following, we obtain a lower bound for *D*_0_. To begin with, for each (*k, 𝓁*) such that *x*_*k*_ ≁ *x*_*𝓁*_, we have the trivial bound

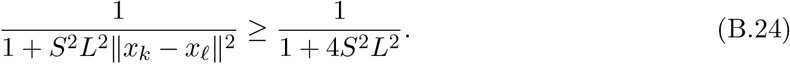

As a result, since

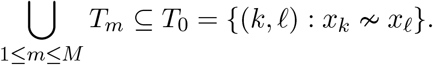

we have

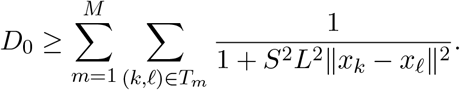

Now since for each *T*_*m*_, if (*k, 𝓁*) ∈ *T*_*m*_ and without loss of generality that *k < 𝓁*, we have

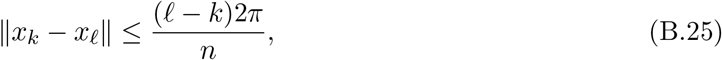

then it follows that

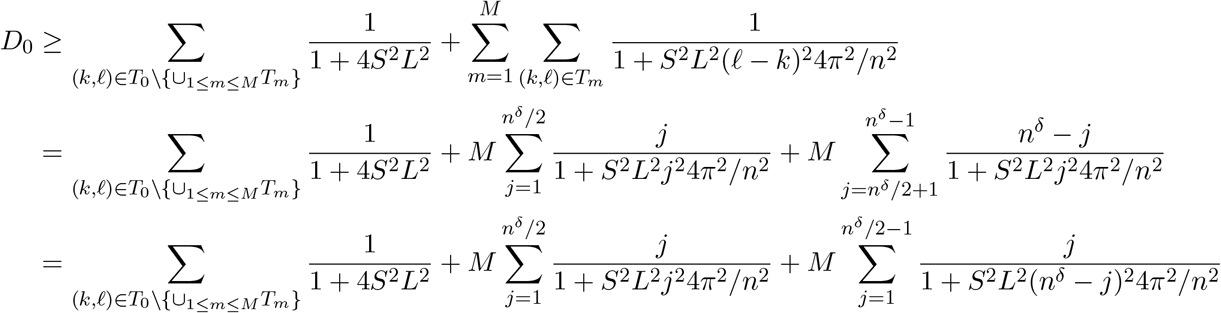

Thus, we have

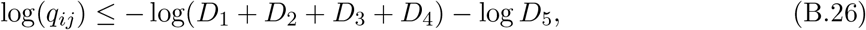

where

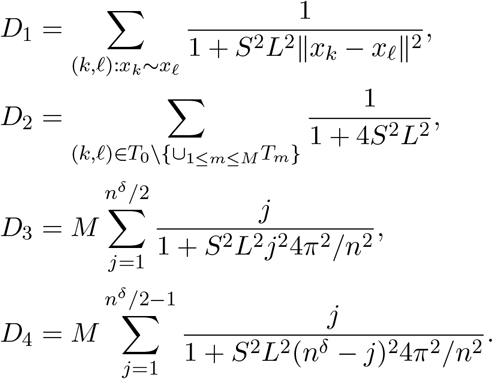

This completes the proof of the theorem

### B.2 Main proof argument

**Proof of (B.1)**. See Proposition B.4. We have

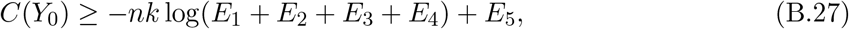

where

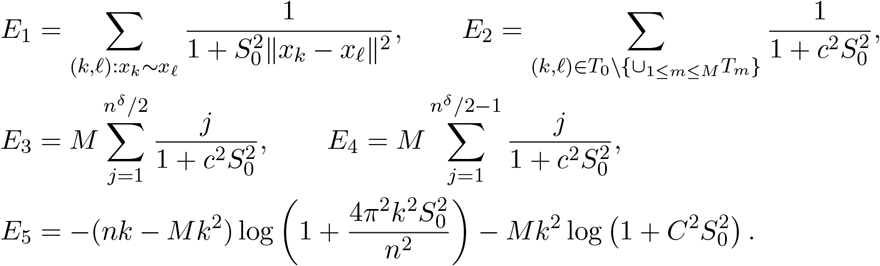

**Proof of (B.2)**. See Proposition B.7. We have

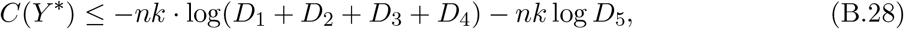

where

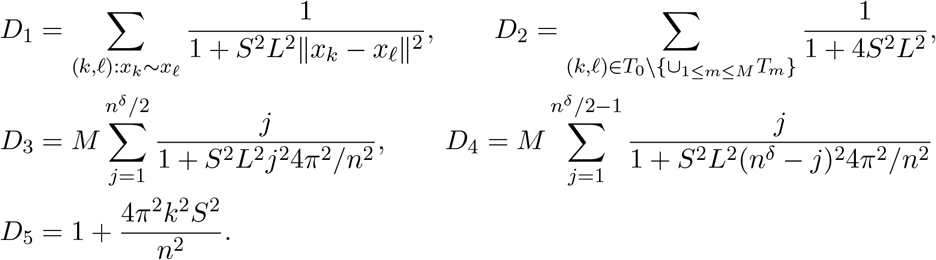

#### Completing the argument

Now by the relation (B.3), we have

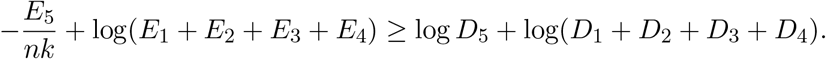

Note that *c* + *δ* ≤ 1 and *m < c*, it then follows that *m* + 2*δ* + *c <* 2*c* + 2*δ* ≤ 2, we have

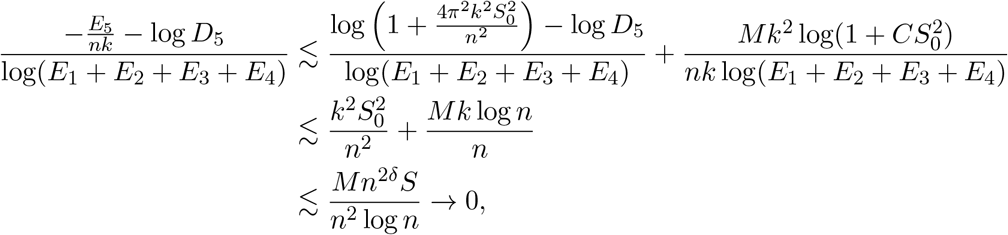

or

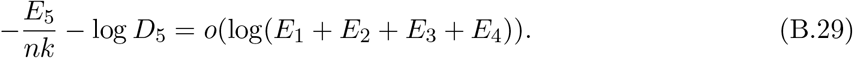

Then it follows that

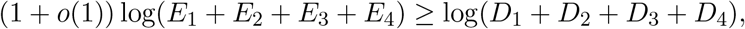

that is, for sufficiently large *n*,

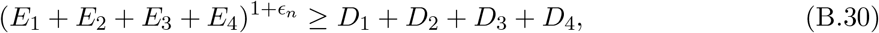

for some series *ϵ*_*n*_ = *o*(1). Note that we can find a sequence of *S*_0_ such that *D*_1_ + *D*_2_ = *E*_1_ + *E*_2_ for all *n*. Specifically, for each *n, D*_1_ + *D*_2_ is monotonic decreasing in *S* and *E*_1_ + *E*_2_ is monotonic decreasing in *S*_0_. For any fixed *S, T* (*S*_0_) = *D*_1_ + *D*_2_ − *E*_1_ − *E*_2_ as a function of *S*_0_ = *θS* satisfies *T* (*S*_0_) *>* 0 if

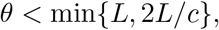

and *T* (*S*_0_) *<* 0 if

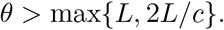

By continuity of *T* (*S*_0_), we know that there exists some *θ* = [min{*L*, 2*L/c*}, max{*L*, 2*L/c*}] (possibly depending on *n*) such that *T* (*S*_0_) = 0. We choose such an *S*_0_ so that *D*_1_ + *D*_2_ = *E*_1_ + *E*_2_. Next, we show that for the above selection of *S*_0_ (so that *S*_0_ ≍ *S*(*ϕ*^*^) or *c* = *τ* ∈ (0, 1)), if in addition *L* ≍ 1, then there exists some series *η*_*n*_ such that *η*_*n*_ ≳ *ϵ*_*n*_ and

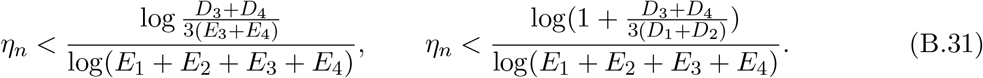

It then follows that

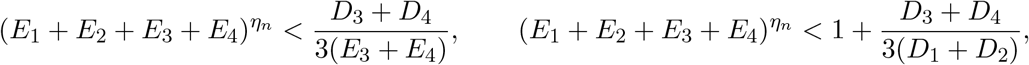

so that

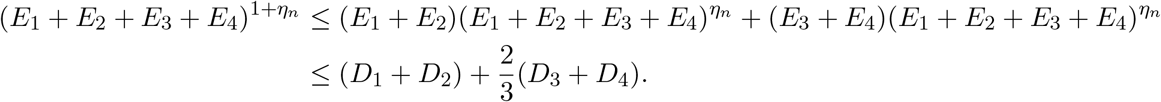

However, this is contradictory to (B.30). Therefore, we conclude that *L* ≍ 1 cannot be true. As a result, we conclude that *L* ≫ 1.

The rest of the proof is devoted to (B.31). By Proposition B.5, we have

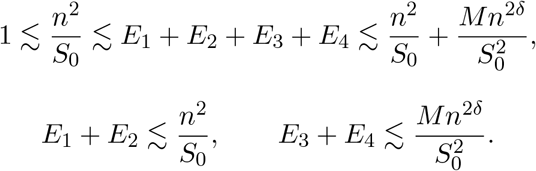

In particular, when

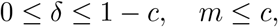

we have

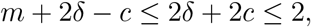

So taht

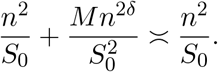

Similarly,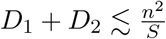. However, for *D*, since *δ* ≤ 1 − *c*, we have

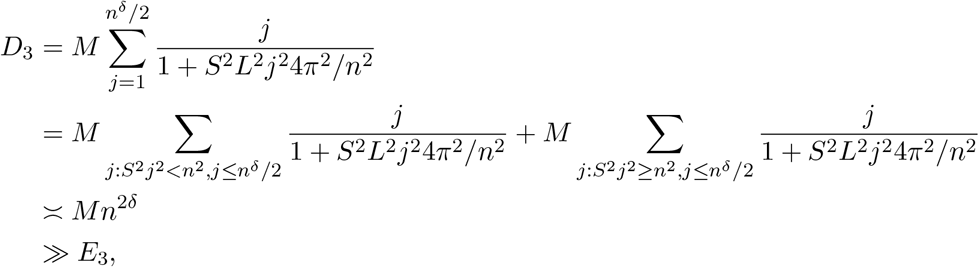

and

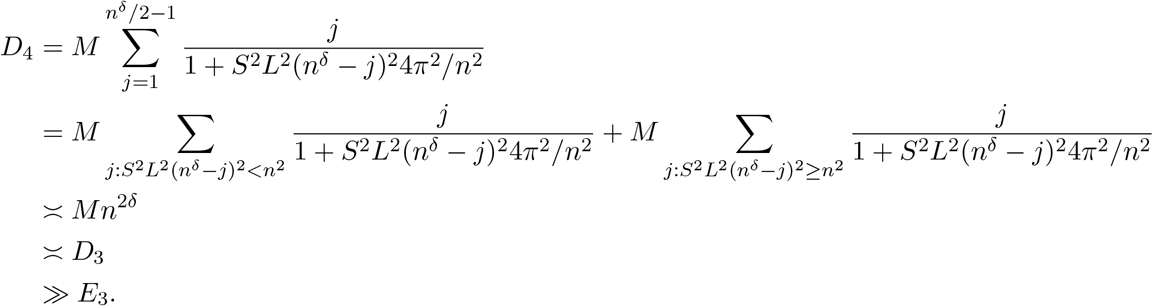

As a result, we have

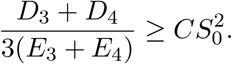

As long as there exists some constant *q >* 0 such that

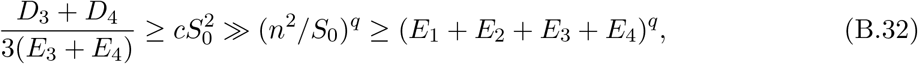

we can choose *ϵ*_*n*_ ≲ *η*_*n*_ *< q* so that the first inequality in (B.31) holds. To see the existence of *q* in (B.32), we can just choose *q* such that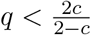 holds. Similarly, since

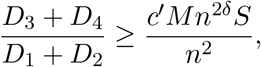

we have

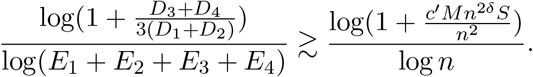

Since

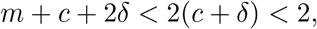

we have

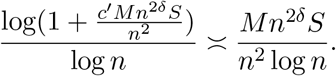

we can choose

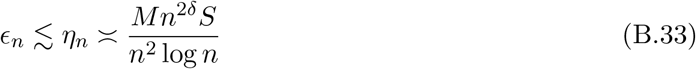

so that the second inequality in (B.31) holds. To see the existence of *η*_*n*_ in (B.33), note that

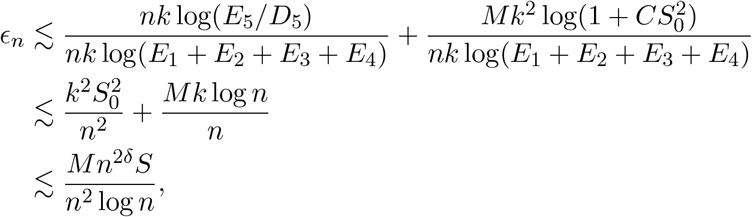

where in the last inequality we used

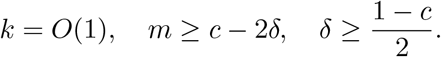

This completes the proof of (B.31).

## Footnotes

1 throughout, we use the words “smooth structure” and “manifold structure” interchangeably.

2 that is, preserving both global and local structures.

3 cell embeddings that do not preserve neighborhood structure as compared with the input data.

